# A transcriptional program shared across lineages underlies cell differentiation during metazoan development

**DOI:** 10.1101/2022.04.22.489139

**Authors:** Marina Ruiz-Romero, Cecilia C. Klein, Sílvia Pérez-Lluch, Amaya Abad, Alessandra Breschi, Roderic Guigó

## Abstract

**Background:** During development, most cells undergo striking changes in order to develop into functional tissues. All along this process, the identity of each tissue arises from the particular combination of regulatory transcription factors that specifically control the expression of relevant genes for growth, pattern formation and differentiation. In this scenario, regulation of gene expression turns out to be essential to determine cell fate and tissue specificity.

**Results:** To characterize the dynamic transcriptional profiles during cellular differentiation, we tracked down the transcriptome of committed cells in different *Drosophila melanogaster* tissues and compartments at a number of developmental stages. We found that during fly development, temporal transcriptional changes shared across lineages are much larger than spatial lineage-specific transcriptional changes, and that cellular differentiation is dominated by a transcriptional program, common to multiple lineages, that governs the transition from undifferentiated to fully differentiated cells independently from the differentiation end point. The program is under weak epigenetic regulation, and it is characterized by downregulation of genes associated with cell cycle, and concomitant activation of genes involved in oxidative metabolism. Largely orthogonal to this program, tissue specific transcriptional programs, defined by a comparatively small number of genes are responsible for lineage specification. Transcriptome comparisons with worm, mouse and human, reveal that this transcriptional differentiation program is broadly conserved within metazoans.

**Conclusions:** Our data provides a novel perspective to metazoan development, and strongly suggest a model, in which the main transcriptional drive during cell type and tissue differentiation is the transition from precursor undifferentiated to terminally differentiated cells, irrespective of cell type.

## Background

All pluricellular organisms develop from a single totipotent cell. In the course of development, cells proliferate and commit to distinct cell fates to ultimately, through cell differentiation, produce a plethora of cell types that combine in specialized tissues and organs. Such a diversity of cell types, all sharing the same genome sequence, is the consequence of differential expression of specific genes, which is driven by complex transcriptional and epigenetic regulatory networks. The conventional view of differentiation explains cell fate commitment as a linear and progressively restricted path that is distinctive for each specific cell type (based on Waddington’s diagram of epigenetic landscape (1)). In this model, the transition from a proliferative state to a differentiated quiescent state is achieved through several cell fate decisions driven by precise epigenetic regulatory programs. However, studies from the last decades in cell reprogramming, transdifferentiation and regeneration have slightly changed this view, by showing that adult cells retain a certain plasticity and that the differentiation process is reversible to varying degrees, both *in vitro*, for example in the case of reprogramming inducible pluripotent stem cells (iPS cells) and *in vivo*, for example in the case of dedifferentiation and transdifferentiation after injury (reviewed in (2)).

In the past two decades, research into the regulatory mechanisms underneath cell fate and tissue differentiation has been enormously facilitated by next generation sequencing (NGS) technologies. Recent single cell sequencing technologies, in particular, have led to the identification of specific gene expression profiles associated with tissues or cell types in adult organisms or after *in vitro* differentiation (3–9).

Here, we analyze the development of *Drosophila melanogaster* --an experimentally manageable model within metazoans-- to characterize the temporal and spatial transcriptional programs that underlie tissue differentiation during animal development. In contrast to previously published fly development transcriptional studies, here we specifically, label primordial cells from imaginal discs --internal epithelial sacs in larvae that are committed to give rise to specific tissues in adults (10)--, isolate them using fluorescence-activating cell sorting (FACS), and profile their transcriptional state with RNA-seq at different developmental stages. Differential gene expression analysis reveals that, in contrast to the prevalent view, a transcriptional program common across cell lineages governs the transition from undifferentiated to fully differentiated cells, dominating tissue specific programs, which are defined by a comparatively low number of genes, and tend to be activated late during development. Comparative transcriptomics analyses show that this program is substantially conserved across metazoans.

## Results

### Spatial and temporal transcriptional analysis of imaginal discs during *Drosophila melanogaster* development

During fly metamorphosis, imaginal tissues undergo cell differentiation and morphogenetic rearrangement to give rise to adult functional appendages (**Fig. 1A)**. To investigate the molecular basis of this process, we interrogated the transcriptome of imaginal tissues at different stages during terminal fly development. Within each tissue and developmental stage, we selected the precursor cells that differentiate into the adult tissue. To track down precursor cells, we used GFP reporter lines, in which GFP is driven by the promoter of genes specifically expressed in a particular region of each tissue. Briefly, imaginal tissues were manually dissected and disaggregated by trypsin treatment. After that, cells were collected by fluorescence-activating cell sorting (FACS), RNA was extracted and processed for NGS (see Methods and **Fig. 1B** and **Supplementary Fig. 1A,B**). Overall we generated RNA-Seq data from eye, leg and wing from three different stages: third instar larvae (L3, around 110 h of development), when cells are predetermined and committed to specific cell types in adult, but they are still undifferentiated and keep proliferative capacity(10)), early pupa (EP, around 120 h of development), immediately after entering pupariation and coinciding with Ecdysone hormone signaling peak, and late pupa (LP, around 192 h of development, corresponding to 72 h after pupa formation), when tissues are fully differentiated and almost functional. In addition, we also generated RNA-seq data for antenna and genitalia discs (male and female), for L3 and EP. Finally, we produced RNA-seq for four wing compartments (anterior, posterior, ventral and dorsal) for the three developmental stages. In total, considering two replicates per condition, we generated RNA-Seq data for 54 samples. In addition, we generated H3K4me3 ChIP-Seq for eye, leg and wing at these three developmental time points (two replicates per condition, 18 ChIP-Seq samples, in total).

**Fig. 1.**
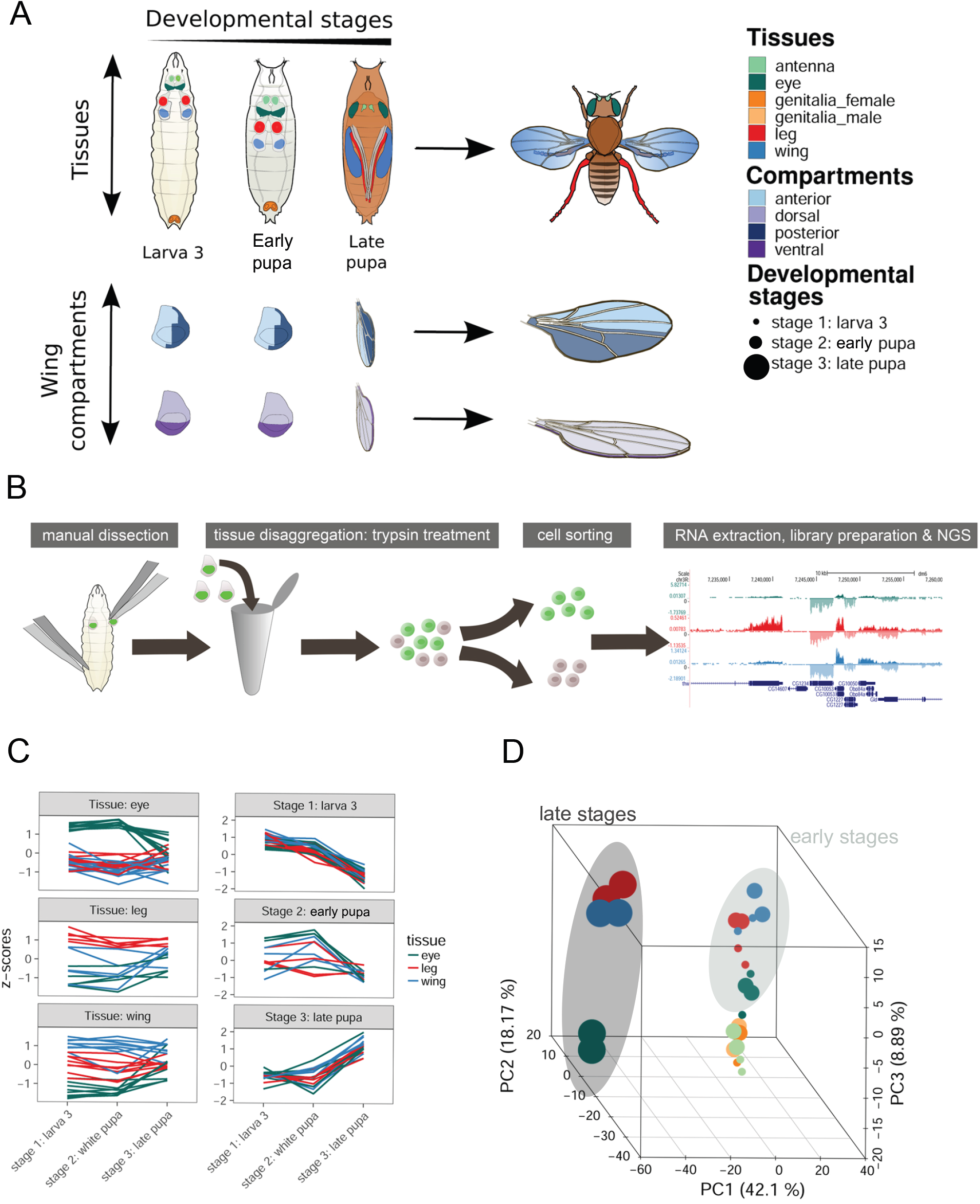
Transcriptional profiling of *Drosophila melanogaster* imaginal discs. **(A)** Overview of tissues, wing compartments and developmental stages profiled along this work. **(B)** Workflow of the RNA-Seq data generation. Briefly, imaginal tissues are manually dissected, disaggregated with trypsin treatment and sorted to collect the cells of interest (i.e., the precursor cells that differentiate into the adult tissues) from each imaginal disc. After that, RNA is extracted and processed for library preparation. **(C)** Expression profiles of genes with known tissue or stage specific expression patterns. **(D)** Principal component analysis (PCA) based on the expression of the 1,000 most variable genes across tissues and developmental stages. Gene expression is computed as log10-normalized Transcripts Per Kilobase Million (TPMs) with pseudocount of 0.01. Only genes with at least 5 TPMs in at least two samples were considered. PC1 separates the early and the late stages. PC2 separates neural and non-neural tissues. PC3 separates the late stage in the eye from the rest of the samples.

From the RNA-Seq data, we estimated expression values for 17,158 annotated genes (FlyBase gene annotation r6.05, summary statistics of RNA-seq samples in Methods, all data available at https://rnamaps.crg.es). Genes with known tissue specific (11–15), or developmental (16–18) transcriptional patterns behaved as expected (**Fig. 1C** and **Supplementary Table 1**).

### A common transcriptional differentiation program is shared across tissues during fly development

Principal component analysis (PCA) and hierarchical clustering (**Fig.1D, Supplementary Fig. 2A, B**) show that samples cluster preferentially by developmental time than by tissue lineage. That is, the transcriptomic profile of a given tissue at a specific stage of development is more similar to the profile of other tissues at that stage, than to the same tissue in other developmental stages. Using linear models (see Methods (19)), we estimated that the proportion of the gene expression variance explained by the developmental time (41% on average) is indeed much larger than that explained by the tissue (22%) (**Fig. 2A, B**). During fly tissue differentiation, therefore, temporal changes of gene expression dominate over spatial changes.

**Fig 2.**
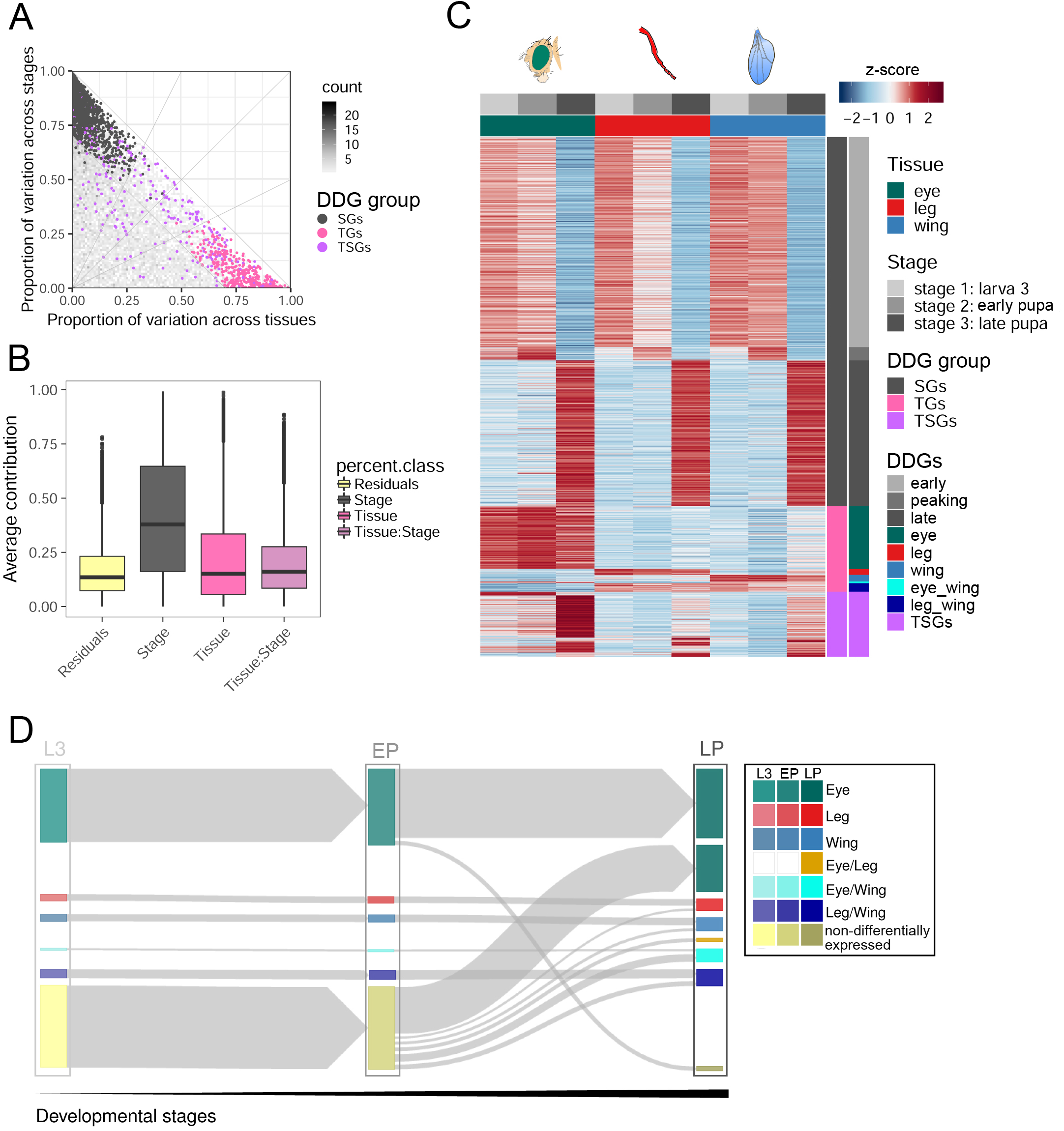
Gene expression dynamics along fly development. **(A)** Proportion of the variance in gene expression explained by tissue (x-axis) and by developmental stage (y-axis). Each dot corresponds to one of the 9,334 genes that are expressed at least 5 TPMs in at least two samples). Genes changing expression across tissues and/or developmental stages (Developmentally Dynamic Genes, DDGs) along fly development are highlighted in grey (differentially expressed across stages, SGs), pink (differentially expressed across tissues, TGs) and purple (differentially expressed across tissues and stages, TSGs). (**B)** Proportion of gene expression variance explained by tissue, stage and the interaction between the two. **(C)** Expression of Developmentally Dynamic Genes (DDGs) along fly development. Gene expression values are normalized to z-score values. **(D)** Dynamics of genes differentially expressed across tissues (TGs) and across tissues and stages (TSGs) represented as a Sankey diagram. At each developmental stage, we represent the genes that are differentially expressed in each tissue and those that are not (yellow). The arrows represent the number of genes that transition from one developmental stage to the next one and from not differentially to differentially expressed in a given tissue (or vice versa). Many tissue specific genes are already differentially expressed at L3 (TGs), but many which are not differentially expressed at this developmental stage become tissue specific at LP (TSGs). There is, in particular, a large expansion of eye specific genes. Overall, the transcriptome diverges as tissues become specified.

From a set of 9,334 genes that are expressed at least 5 TPMs in at least two samples, we identified a set of 2,034 genes that change expression across tissues (eye, leg and wing) and/or time points (Developmentally Dynamic Genes, DDGs, see Methods(20)). We classified these genes in three categories: differentially expressed across developmental stages (stage genes, SGs, 1,445 genes, consistently with the result above, the largest category), across tissues (tissue genes, TGs, 345) and across both tissues and stages (tissue-stage genes, TSGs, 255), **Supplementary Table 2**). Within SGs, we further classified genes as downregulated or early differentiation genes (822 genes, 56%, preferentially expressed in L3 and EP), upregulated or late differentiation genes (571 genes, 39.5%, preferentially expressed in LP), and peaking or metamorphosis entrance genes (52, 4.5%), which are upregulated at EP in all tissues (**Fig. 2C** and **Supplementary Fig. 3A,B**). Early genes are associated with cell cycle, gene regulation, RNA processing and translation processes; metamorphosis entrance genes to endoplasmic reticulum localization and apoptosis signaling, and late genes are related to cuticle formation and chitin metabolism (**Supplementary Fig. 3C**). Among the late genes, a large fraction (324, 57%) are poorly characterized genes with no associated functions, compared with 42% of metamorphosis entrance genes, and 27% of early genes. This could be ascertainment bias, as most functional characterization studies in flies are performed in developing, not in fully differentiated, animals.

TGs are enriched for the expected tissue-specific cell fate functional categories (**Supplementary Fig. 3D**). Some of these genes are known to be essential to regulate cell determination and tissue formation during development (e.g. *ey* and *gl* in eye). TGs correspond broadly to genes that are already differentially expressed at L3, when cells within imaginal tissues are undifferentiated, and remain differentially expressed all through development. TSGs, in contrast, are genes activated in a tissue specific manner only at specific developmental time points. We found that most TSGs are specifically activated in the transition from EP to LP, driven mostly by an expansion of eye specific genes. This results in a larger number of tissue specific genes at the terminal stage of differentiation associated to each tissue function (**Fig. 2D and Supplementary Fig. 3E**). Transcriptional differences between tissues, therefore, increase with developmental time (**Fig. 1D, 2D**), suggesting that an expansion of tissue regulatory programs is needed for terminal tissue differentiation.

Overall, our results strongly suggest that during fly development, there is a temporal transcriptional program common to all tissues that dominates over tissue specific transcriptional programs, which are defined by a comparatively small number of genes. This program is of fractal nature, as it can be observed at different organizational scales. Indeed, we produced and analyzed expression data from the distinct compartments within wing imaginal discs. As with tissues, variation of gene expression is much larger across developmental time than across compartments (**Supplementary Fig. 4A,B,C**) and the transcriptional behavior of DDGs within the wing imaginal discs replicates the behavior observed among imaginal discs during development (**Supplementary Fig. 4D**).

We have investigated the epigenetic features underlying the fly temporal differentiation program. We generated H3K4me3 ChIP-Seq profiles for eye, leg and wing differentiating tissues and analyzed FAIRE-Seq data available for these tissues to assess chromatin accessibility (21). We focused on the core promoters (+-250bp from the transcription start sites, TSS). Overall, we found most DDGs either in closed conformation and/or unmarked, or unspecifically open and/or marked (**Fig. 3A, Supplementary Fig. 5A-D**), Chromatin accessibility, and H3K4me3 marking in particular, mostly reflect, actually, the breadth of gene expression. Genes with restricted expression (expression restricted to a single stage or/and tissue) are in closed chromatin conformation or/and unmarked more often than genes with widespread expression (**Fig. 3A, Supplementary Fig. 5A-E)**. This is consistent with previous reports that show absence of marking by canonically activating histone modifications in genes regulated during fly development (15,22,23). The exception are early genes with restricted expression patterns. These tend to be marked at early developmental stages and remain marked in late differentiation, even when not being expressed.

**Fig. 3.**
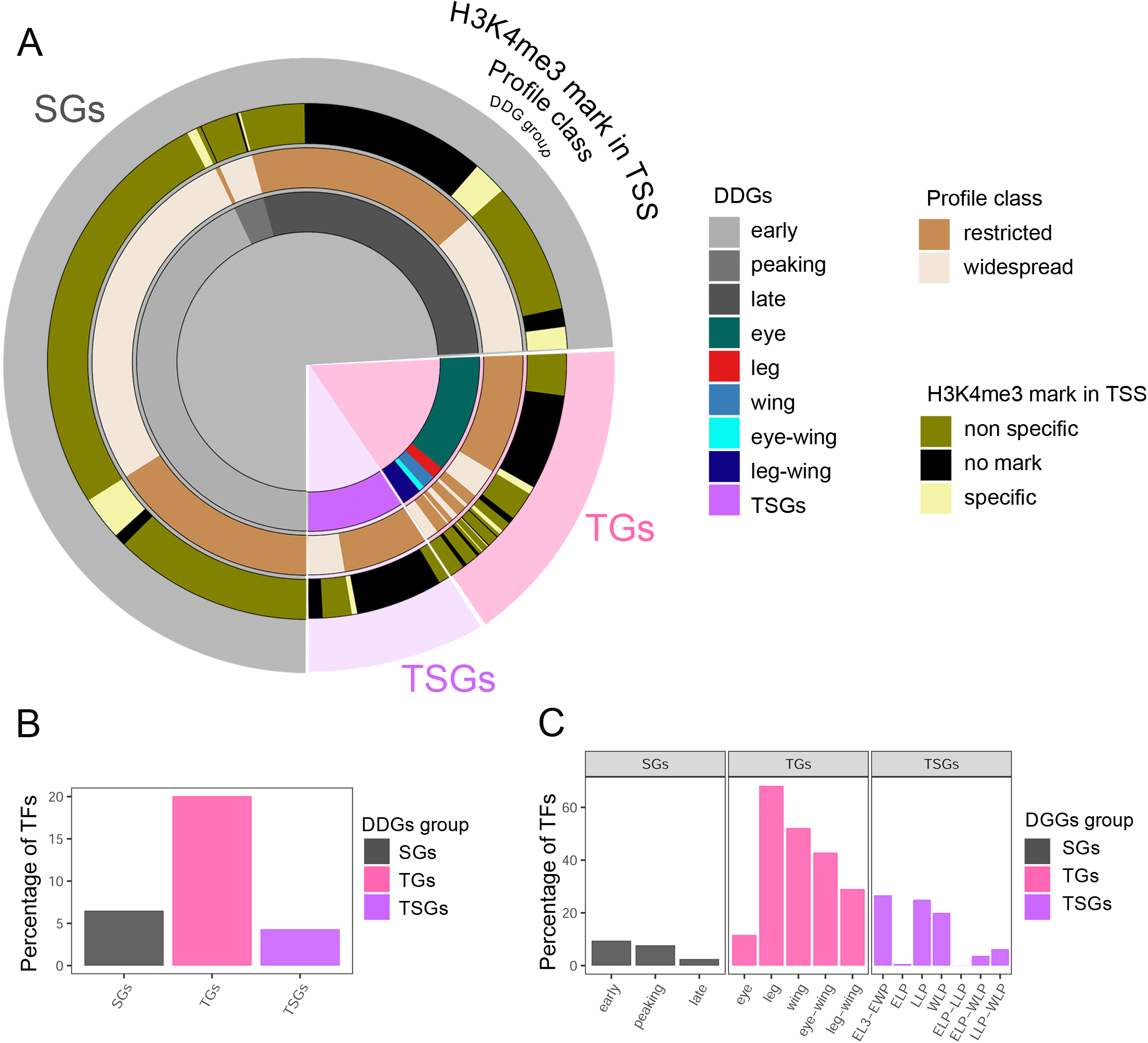
Regulation of DDGs. **(A)** Epigenetic regulation of DDGs. The innermost circle labels genes according to DDGs classification; the second circle displays the breadth of gene expression (profile class), and the third circle displays H3K4me3 marking. **(B** and **C)** Percentage of TFs in DDGs categories.

These results suggest that the fly transcriptional developmental program is, broadly, under weak promoter epigenetic regulation. We did find, however, a strong enrichment of transcription factors (TFs) within TGs (20%) compared to SGs (6%) or TSGs (5%, Fisher’s Exact Test on Count Data, p<0.001) (**Fig. 3B,C)**, suggesting that TFs play a comparatively more important role in spatial than in temporal differentiation. Within SGs the number of TFs is higher in early than in late genes, in concordance with previous observations during mammalian development (24).

### A developmental gene regulatory network in fly differentiation

Gene regulatory networks (GRNs) integrate information from transcriptional regulators and their targets to provide a holistic view of the regulatory program of a particular biological process. Previous studies have successfully used transcriptional-based GRN to model gene regulation along developmental processes (12,25,26). Here, we used Weighted Correlation Network Analysis (WGCNA) (27) to construct a differentiation co-expression network connecting Developmentally Dynamic genes (DDGs) with their putative regulatory TFs. To identify reliable TF-target pairs, we scanned for conserved TF binding motifs occurring in open chromatin regions within the promoter regions of the target genes (see Methods for details, **Supplementary Fig. 6A**).

The resulting GRN includes 1,656 nodes (1,485 DDGs and 229 TFs), and 14,039 edges (**Fig. 4A, Supplementary Fig. 6B-F, Supplementary Fig. 7 A,B** and **Supplementary Table 3**). The WGNA identified 15 regulatory clusters (**Fig. 4B, Supplementary Fig. 6G-I**), to which we associated functional categories by GO enrichment analysis (**Fig. 4C, Supplementary Fig. 8** and **Supplementary Table 3)**. Clusters 1 to 3 form a super-cluster mostly composed of early genes. Cluster 1 is functionally associated with regulation of gene expression (**Fig. 4C, Supplementary Fig. 8**) and, as expected is the cluster with the highest number of interactions (8,486, **Fig. 4D**). The TFs in the cluster include general transcriptional regulators, known to have functions on chromatin structure and regulation of gene expression like BEAF-32, Dref, Z, Dalao, BAP170 or Br; developmental chromatin remodelers such Trl, Pcl, Pho and Phol; insulators like Cp190 and SuHw; TFs associated to imaginal disc morphogenesis, like Hth, Exd, Sd, Da, Mad, Ets21C, Rn, Myc or Max; as well as TFs related to metabolic regulation, like Bigmax, Foxo and ATF-2 (**Supplementary Fig. 7C**). Although these TFs bind many early genes, interactions with late genes and TGs were also predicted. Clusters 2 and 3 are functionally associated with cell cycle and translation, respectively. Clusters 4 and 5, corresponding to very few peaking genes (**Supplementary Fig. 6H**) could also be included in this super-cluster (**Fig. 4A**).

**Fig. 4.**
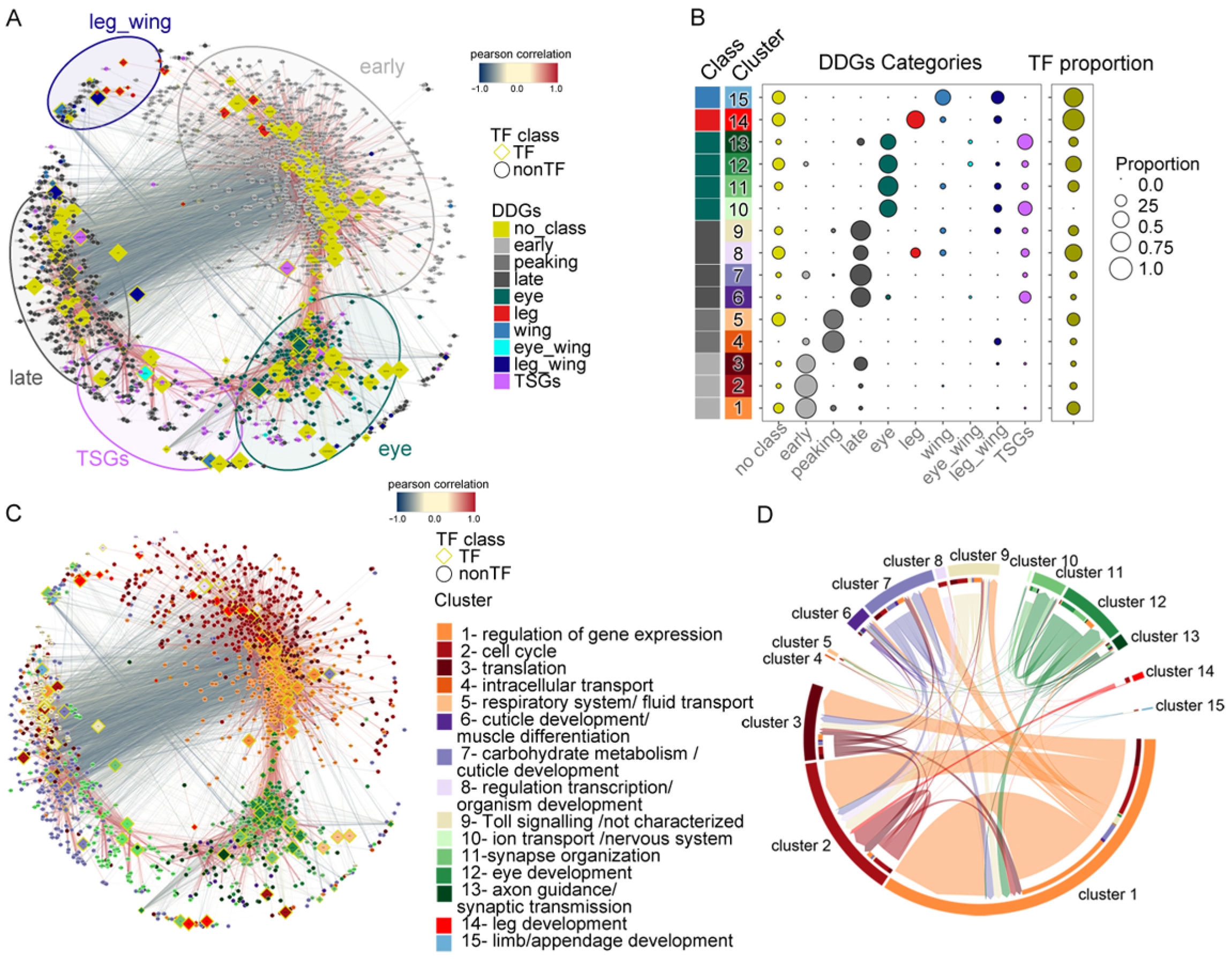
*Drosophila* gene regulatory network (GRN). **(A)** GRN. Edges are colored according to TF-target Pearson’s correlation coefficient (red > 0.3, blue < -0.3). While most correlations are positive, negative interactions are predicted between early and late gene clusters as well as between clusters corresponding to different tissues. Node size reflects node closeness centrality (that is, how close a node is to all other nodes). Nodes are colored according to DDG classification. **(B)** Network clusters. Proportion of DDG categories and TFs in each cluster. The class column indicates the most abundant DDGs category within each cluster. **(C)** GRN. Nodes are colored according to the clusters to which the genes belong and the GO categories associated. **(D)** Connectivity between clusters. Arrows indicate directionality from TFs to targets. The width of the arrows is proportional to the number of TF-target pairs. For ease in interpretation, we have included inner colored bars, representing the target’s clusters.

Clusters 6 to 9 form a second super-cluster mostly composed of late genes. Only one of these, cluster 6, is preferentially regulating late genes, while the TFs within cluster 7 and 9 are predicted to have high number of interactions also with early genes, suggesting potential negative regulation (**Fig. 4C,D)**. Among these TFs, many are activated through stress, immune and hormonal signaling pathways, like dl, Gce, Hr4, Kay, Rel, Eip74EF, Eip75B and Eip93F; and many are known or predicted to have repressor capacity. This hints at a possible link between whole animal signaling and SGs. Signaling cascades, likely associated with metamorphosis, could induce expression of TFs that may repress early genes and activate late genes. Additionally, cluster 8 includes ten TFs: Abd-B, Awh, D, Gsb, Gsb-n, Lim1, Odd, Opa, Retn, Sob, all known to play a role in development and patterning, that predominantly interact with cell cycle genes (cluster 2, **Fig. 4D**).

Clusters 10 to 13 form a third super-cluster mainly composed of eye genes and eye late genes. They are functionally associated with eye morphogenesis and neural fate. Among them, cluster 12 is associated specifically with eye development, and includes eye fate regulators and neurogenesis inductors predicted to preferentially bind eye genes (**Fig. 4D**). These include well characterized genes like: *ey, gl, mirr, toy, oc, pnt, ro, scrt, ttk, lola* and *so* (**Supplementary Fig. 7D**). Some TFs inside these clusters, which are not necessarily expressed in a restricted manner in the eye, are predicted to also bind several early genes. The same is observed for TFs in cluster 14 (leg) and 15 (leg and/or wing, **Fig. 4D**), which form the fourth super-cluster. These results point to a possible crosstalk in which tissue fate regulators mediate down-regulation of early genes when tissues differentiate.

TF-target interactions tend to take place within super-clusters that are associated to different temporal (early, late) and spatial (eye, leg-wing) niches (**Fig. 4D**), suggesting that the temporal expression program is largely orthogonal, with some, but little, cross-talk with tissue specific regulatory programs (**Fig. 4A,C,D**). Further plotting the network according to betweenness centrality, in which highly connected nodes are placed in proximity irrespective of the direction of the correlation (**Supplementary Fig.7A,B**), revealed that early genes are highly connected with TFs regulating a large number of genes, while late genes are more sparsely connected to their regulators, consistent with our observation that tissue specific expression programs unfold late during development (**Fig. 1C, 2C,D**).

The GRN helps to functionally characterize the temporal differentiation program shared across tissues. In this program, the transition from precursor to differentiated states is driven, independent of cell type, by downregulation of genes associated with cell cycle (cluster 2), with regulation of gene expression (cluster 1) and with translation (cluster 3), and by a concomitant activation of genes involved in oxidative metabolism and other metabolic pathways (cluster 7), and in (terminal) differentiation (cluster 6 and 8).

### The fly transcriptional differentiation program captures the transition from undifferentiated to terminally differentiated cells, irrespective of the differentiation end point

To further corroborate our results, we have analyzed other fly developmental transcriptome data, which is not from tissue isolated cells, but from the whole body and from specific tissues. First, when analyzing data from carcass, central nervous system (CNS) and fat body available for L3 and LP or adult(22,28), we found that the expression of SGs clearly differentiates early from late stages (**Fig. 5A, Supplementary Fig. 9A**). This was true in particular for carcass and CNS that, as imaginal discs, contain mostly undifferentiated cells in L3 and experience differentiation during metamorphosis, while the fat body is already differentiated at L3 (review in (29)).

**Figure 5.**
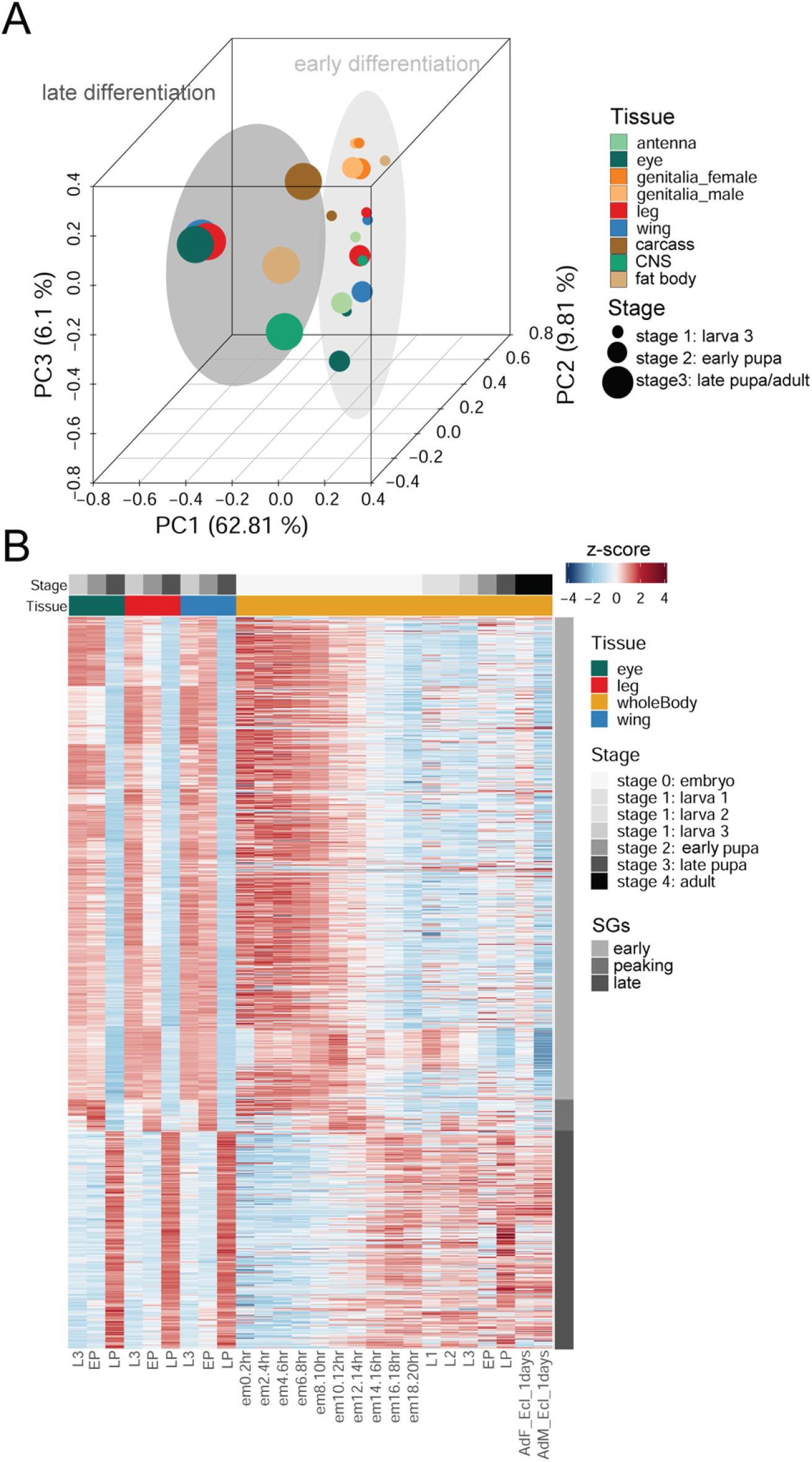
Gene expression dynamics of SGs in tissues and whole animals during fly development. **(A)** Principal component analysis (PCA) of SGs based on modENCODE tissue data (CNS and fat body from L3 and LP and carcass from L3 and adult) and the imaginal tissue data produced here at L3, EP and LP. **(B)** Expression of SGs in modENCODE whole animals from embryo to adult stages, and in the imaginal tissues monitored here at L3, EP and LP.

Next, we have analyzed whole body RNA-seq data from the modENCODE project which is available at much higher temporal resolution (28,30)) (**Figure 5B**). We found that early differentiation genes are highly upregulated at the beginning of embryogenesis, coinciding with active proliferation state, and their expression decreases around mid embryogenesis, coinciding with morphogenetic arrangements and organ primordia specification. On the contrary, late differentiation genes appear upregulated from mid to late embryogenesis, and in larval and pupa stages compared to early embryogenesis. As expected, when measured on the whole organism, containing heterogeneous cell types in different states of differentiation, the shift between early and late gene expression can not be detected comparing L3 and LP stages (**Fig. 5B** and **Supplementary Fig 9B**). This suggests that the fly developmental transcriptional program is actually associated with cell differentiation, and that the endpoints of this program (early and late genes) correspond to undifferentiated and fully differentiated cells, rather than to specific chronological differentiation time points. In additional support of this, we have analyzed RNA-Seq data available for different cell types from the *Drosophila* adult midgut (31). We found that in undifferentiated or primordia cells (intestinal stem cells and enteroblasts, respectively) early genes are up-regulated compared to differentiated cells (enterocytes and enteroendocrine cells), in which late genes are up-regulated, instead (**Fig. Supplementary 9C**).

### The fly transcriptional differentiation program is conserved in metazoans

To investigate whether the fly temporal differentiation program is conserved outside from insects, we analyzed RNA-Seq data from diverse organs at different developmental time points available for mouse, human, and worm (24,32). We identified the 1-to-many orthologs of the set of fly early and late genes (**Supplementary Table 4**) in each of these species. In the case of mouse, more than 80% of orthologs of fly early and late genes were classified identically by Cardoso-Moreira et al. (24) in at least one of the mouse tissues (**Fig. 6A**). Consistent with tissue specialization during development, also observed in *Drosophila*, while 38% of early orthologs are downregulated through differentiation in all four tissues (325 out of 850), only 11% of late orthologs are upregulated in the four tissues (42 out of 372, **Fig. 6A**). In agreement with the metabolic changes observed during fly development, early and late mouse orthologs are functionally associated to different metabolic pathways, including nucleic acid metabolism for early genes orthologs and ion transport and lipid metabolism for late ones (**Fig. 6B**).

**Fig. 6.**
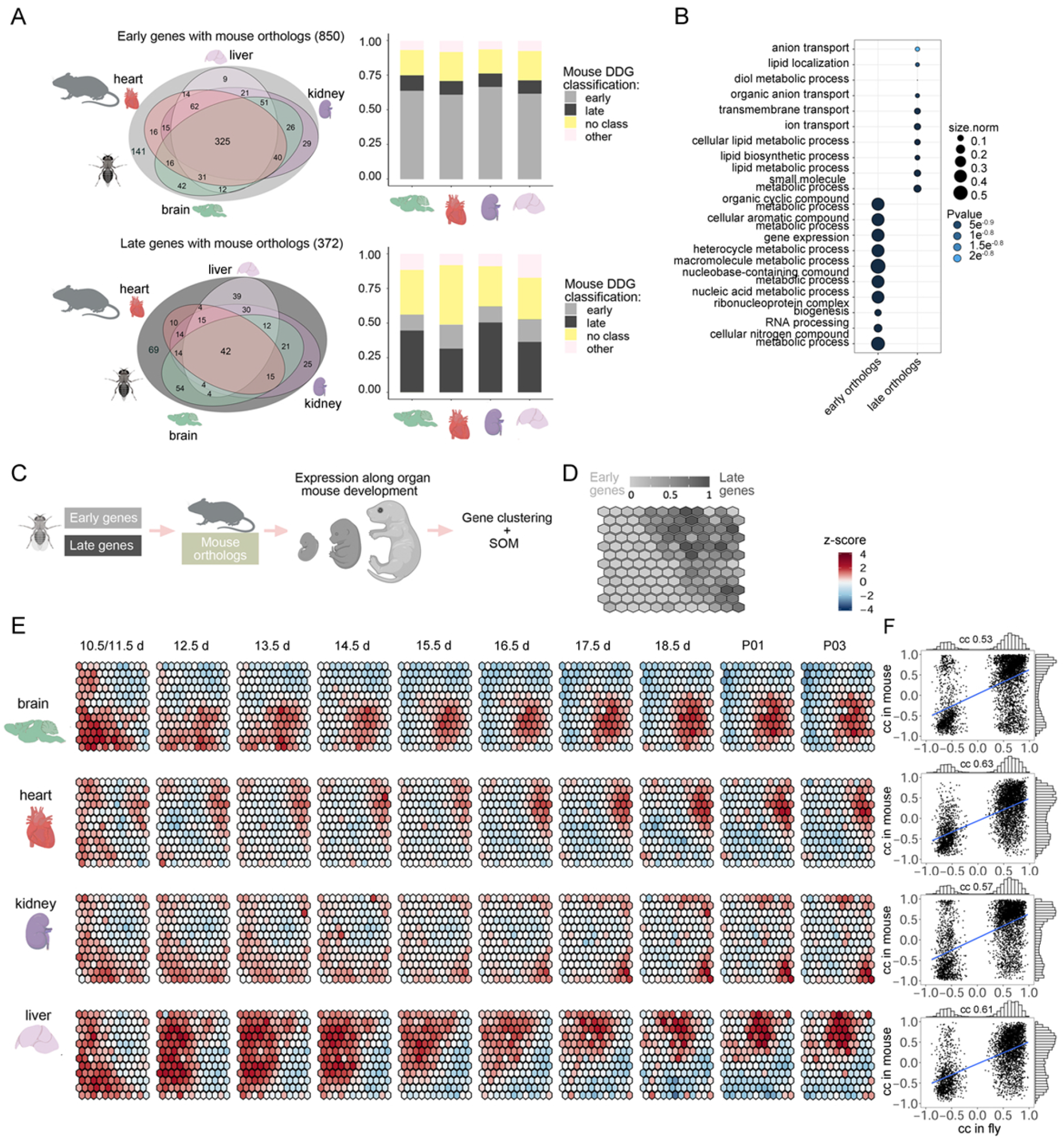
Conservation of the *Drosophila* GRN in mouse. **(A)** Venn diagrams (left panels) showing the number of fly early and late genes, the orthologs of which are also classified as early and late during mouse development according to Cardoso-Moreira et al. (*24*) The bar plots (right panels) show the early and late orthologs classified by expression profile in mouse tissues according to Cardoso-Moreira et al.(*24*) **(B)** GO term enrichment analysis of early and late orthologs in mouse. **(C)** Mouse orthologs of fly early and late genes are clustered using self-organizing maps (SOM) based on the RNA-seq derived expression from a number of organs during mouse development (brain, heart, kidney and liver from embryo 10.5/11.5 days through 3 days post-natal stage). **(D)** SOM clustering of mouse orthologs of fly early and late genes. Each cell corresponds to a gene cluster. We are considering 12×12 cells, each containing between 2 and 23 genes. The grey intensity of cells denotes the proportion of early (light grey) versus late (dark grey) genes in the cell/cluster. Early and late orthologs cluster separately regarding gene expression along mouse organ development. Early genes cluster preferentially on the left part of the SOM representation, while late genes cluster on the right. **(E)** Changes in expression along mouse development in brain, heart, kidney and liver of early and late orthologs. **(F)** Scatter plots of the correlation in fly and in mouse of TF-target pair. The correlations have been computed independently in each fly-mouse ortholog tissue. When multiple mouse orthologous targets are found for the same orthologous TF, the TF-target pair with the closest correlation is employed. Pearson’s correlation coefficient (cc) between fly and mouse correlations is shown on the top of the plot, p-values of the correlations are 1.09^e-317^ for brain and 0 for heart, kidney and liver. (Figure partially created with BioRender.com).

We used self-organizing maps (SOM) to cluster the orthologous genes in each species based on the developmental gene expression data in that species (**Fig. 6C-E and Supplementary Fig. 10**). In the case of mouse, orthologs of fly early and late genes (corresponding to 850 and 372 fly genes, respectively) clearly cluster apart (**Fig. 6D**). In every tissue analyzed, the gene expression trajectory during differentiation followed a similar path, replicating that observed in the fly, with higher expression of early orthologs in the first time points of development gradually transitioning to higher expression of late orthologs in the last time points (**Fig. 6E**). While early orthologs have similar widely distributed expression patterns in early development, late orthologs show tissue-specific specialization late in development.

Next, we further investigated whether the specific associations TF-target detected in the fly GRN were conserved in the mouse. We computed the correlation of expression between orthologous TFs and orthologous targets across mouse samples for each tissue separately. Since for each fly TF-target pair there may be multiple mouse orthologs TF-target pairs, we selected the mouse TF-target pair with the closest correlation to the fly TF-target pair, irrespective of the direction of the correlation. In all tissues, we found the TF-target associations (direction and strength) strongly correlated between fly and mouse (**Fig. 6F**). We believe that the assumption that the best correlated pair is the one most likely to have kept the fly function in the mouse after the subfunctionalization and neofunctionalization, expected to occur following gene duplication, is the most sensible one. However, it may also lead to inflated correlation values. Thus, we have recomputed the correlations when considering all orthologous pairs for each fly TF-target pair. While the correlations are, as expected, weaker, they are still highly significant (**Supplementary Fig. 11**).

We found similar results when analyzing human and worm developmental expression data (**Supplementary Fig. 10**). We then identified the most conserved TF-target pairs: 77 pairs in the fly (nine TFs, 68 targets, **Supplementary Table 5**) having correlations higher than 0.5 in all species). Among these, the most conserved pairs include the Hox cofactor Exd and the TGF-Beta related factors Mad and Med as TFs. These TFs are predicted to regulate several transcriptional and chromatin factors across all metazoans, like the members of the Brahma complex: *Brm, Bap60* and *Dalao, Row, Glo* and *CG1620*, some splicing factors like Hel25E and B52 and some cell cycle regulators like Mapmodulin and Grp. Also Max, the cofactor of Myc, is in the list of most conserved pairs regulating *RpS30*, involved in translation, and *CG8209* (the ortholog of *UBXN1*, a general negative regulator of protein metabolism.

These results, all together, strongly suggest that the fly transcriptional differentiation program is largely conserved within the metazoan lineage.

## Discussion

Cell determination and differentiation are fundamental to tissue and organ formation and ultimately organism development. To characterize the molecular basis of these processes, we profiled gene expression of imaginal tissues during fly organ differentiation. In contrast to previous work (4,18,21,33,34), we specifically labeled primordial cells with GFP and tracked them along development. This allowed us to monitor changes precisely associated to particular cells while they undergo differentiation (**Fig 1A and Supplementary Fig. 1**). The data produced here, therefore, is a valuable resource to investigate transcriptomic changes in fly development.

Our analyses of this data suggest that the transition from precursor undifferentiated to terminally differentiated cells is the consequence of two, partially orthogonal, transcriptional programs. First, the general down-regulation of early genes and activation of late genes, which is common to all differentiating cells, independently from the differentiation end point. Second, the late specialized activation of genes defining tissue fate (**Fig. 7A**). The temporal differentiation program clearly dominates the spatial program, which is defined by a relatively small number of genes. This suggests that the Waddington landscape is less steep, and the valleys less deep than often assumed, contributing to explain why transdifferentiation from a terminal cell type to another can be forced with relative ease, either directly or through de-differentiation.

**Fig. 7.**
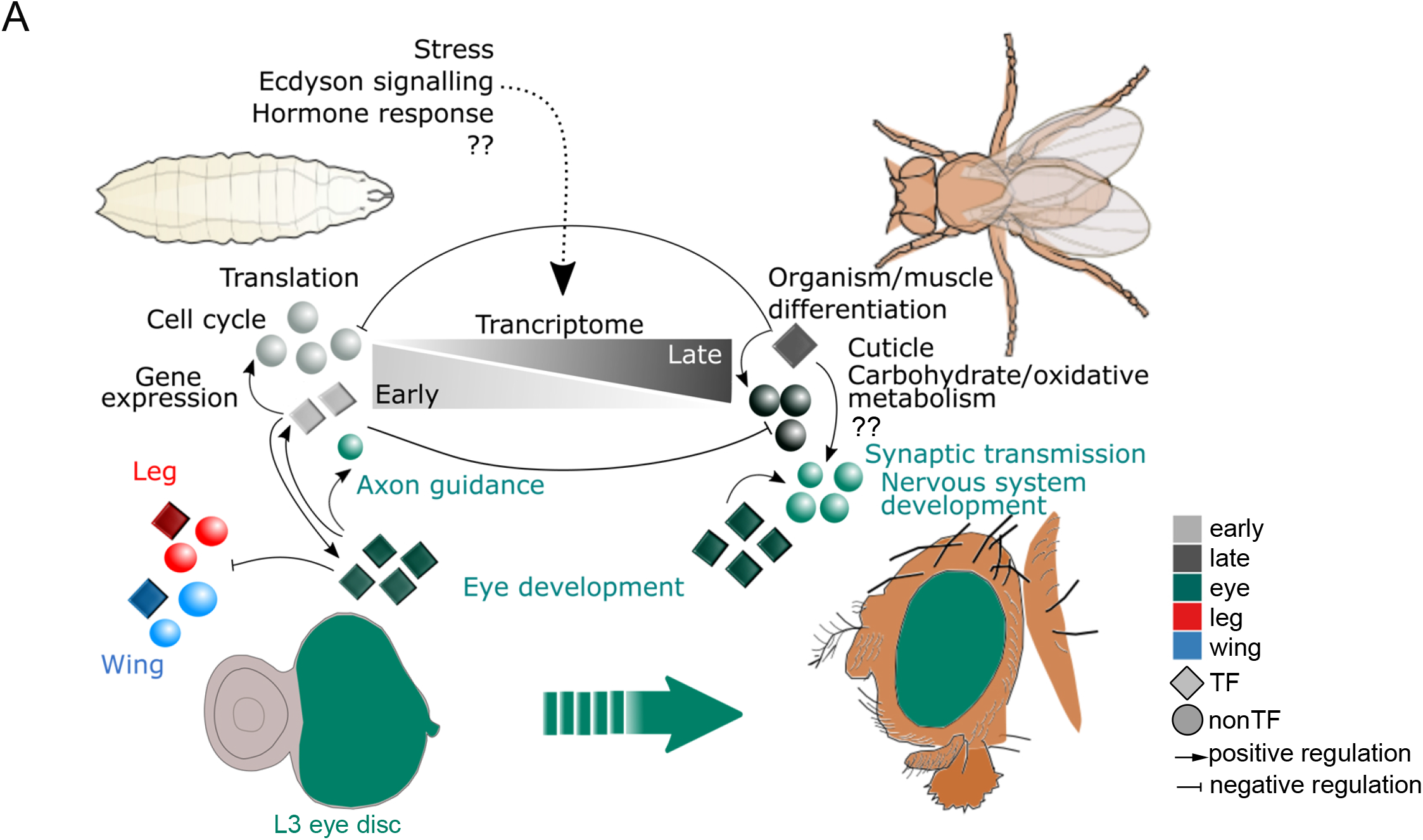
Model of gene regulation in *Drosophila* differentiation. **(A)** General view of tissue differentiation (e.g. eye differentiation, green). Differentiation requires a transcriptional switch from early genes (light grey) towards late genes (dark grey) that could be mediated by stress, hormonal signals or other uncharacterized systemic and external signals (??). Gene expression, translation and cell cycle genes decrease while cuticle, organism differentiation, oxidative metabolic genes and functionally uncharacterized genes (??) are up-regulated upon differentiation. Eye genes, mainly involved in cell fate regulation, are expressed from precursor cells to fully differentiated ones along the process. Eye specific TFs regulate eye genes as well as TSGs. Early eye genes (green) are associated with axon guidance function and late eye genes are related to synaptic transmission and nervous system development functions. Negative regulation occurs between early and late genes and among tissue-specific TFs.

We have built a differentiation gene regulatory network (GRN, **Fig. 4**) to help characterize these programs from the functional standpoint. We found that genes generically downregulated during differentiation are preferentially associated with cell cycle, regulation of gene expression and translation. Remarkably, most genes (∼ 60%) generically up-regulated during differentiation have not been functionally characterized yet. We identified a cluster, however, in the fly differentiation GRN (cluster 7), mainly composed of late genes involved in oxidative metabolism and ion transport. These include *Vha14-1, Vha36-1, Vha68-2, VhaAC45 and VhaM9.7* (35) that encode for subunits of vacuolar H+ ATPase; and genes related to other metabolic pathways, like *mmy* (36), *Gs1* (37), *Mfe2* (38) among others (**Fig. 4 and Supplementary Fig. 7**). White et al. (39) previously described a transcriptional metabolic change associated with metamorphosis entrance in *Drosophila*, and hypothesized that this could reflect tissues preparing for cell death, or consequence of the transition from an active larval state to a sessile pupa one. However, recent insights in early embryogenesis and stem cell (SC) differentiation (reviewed in (40)) indicate that in mammalian embryonic SCs (ESCs) oxidative capacity is reduced and glycolysis-dependent anabolic pathways are enriched whereas mitochondrial function and oxidative metabolism positively correlate with SC differentiation. Thus, experimental evidences based on mammalian ESCs reprogramming and differentiation indicate that transcriptional programs regulating stemness influence energy metabolism and metabolic enzymes ((41–44) review in (40)). In agreement with this, we found that mouse orthologues of the early and late gene sets are enriched for genes associated with different metabolic pathways (**Fig. 6B**). Altogether, our results suggest that the fly transcriptional differentiation program may be regulating metabolic changes necessary for tissue differentiation (**Fig. 7A**). Metabolomic assays to characterize the metabolic fluxes occurring in imaginal tissues through differentiation would help to assess this hypothesis.

The fly transcriptional differentiation program, thus, appears to be under weak direct epigenetic regulation. Chromatin accessibility appears quite stable and unspecific for genes regulated during fly development, according to previous publication (21) (**Supplementary Fig. 5C,E)**, and it does not seem therefore to play a key direct regulatory role. However, it could still play an indirect role through the regulation of certain key fate regulators in specific tissues. For example, we found three eye TFs (Scrt, Oli and Hmx) showing open promoters specifically in the eye. These genes are involved in eye and neural fate specification, and regulate multiple genes according to our GRN. For instance, among Scrt direct targets there are essential eye fate regulators like Ey, So and Pnt. Marking by H3K4me3, on the other hand, reflects mostly the breadth, rather than the specificity, of gene expression **(Fig. 3, Supplementary Fig. 5**). We specifically found, however, that genes exclusively expressed in L3, tend to maintain the mark in LP. H3K4me3, thus, is not actively erased from the switched-off promoters of these genes. While the consensus in the field is that H3K4me3 is a conserved hallmark of active promoters, recent reports (reviewed in (45)) show that it is dispensable for gene activation; our results further suggest that it may not be sufficient either to drive and/or maintain transcription.

In contrast, the fly GRN predicts regulatory TF-target interactions for 74% of the genes regulated during differentiation (DDGs). This suggests that the combinatorial action of TFs on open promoters are likely to play the leading role in the regulation of DDGs during cellular differentiation (reviewed in (46,47)). Further studies on TF occupancy in promoters and enhancers during imaginal tissues differentiation, as well as direct and indirect protein-protein interactions between TFs, are required to fully understand the regulation of the transcriptional output of DDGs.

We found that the fly transcriptional differentiation program is broadly conserved among metazoans (from worm to humans) (**Fig. 6E,D and Supplementary Fig.10**). Specifically in the fly, we found that temporal transcriptional changes common to multiple cell types, as they transition from undifferentiated to fully differentiated types, dominate over cell types specific transcriptional changes. It has been recently shown that also during mammalian organ development there is a temporal transcriptional program common across tissues (24,26). However, in this case, tissue specific transcriptional changes seem to predominate (24,26). This does not necessarily reflect a true biological phenomenon, but could partially be the consequence of the difficulties of measuring in systems of large complexity. First, mammalian organs are composed of a large number of cell types and tissues, which do not necessarily differentiate and mature in a synchronous manner, especially in late development. In contrast, the fly organs, at least those analyzed here, (eye, leg and wing), are simpler, composed by a reduced number of cell types (48–50), which, as a result of metamorphosis, mature in a more synchronized manner. Second, as vertebrates suffered several rounds of whole genome duplications, duplicated genes underwent different paths of neo-and sub-functionalization (51,52), evolving divergent(53–55) or redundant regulatory profiles (56,57) and making difficult to correctly identify TF-target interactions. In *Drosophila*, in contrast, reduced genome complexity facilitates the identification of these interactions. This can actually be seen when comparing the fly TF-target interactions in other species. Thus, in worm, with reduced genome complexity compared to human and mouse, the drop in TF-target correlations when computed over all orthologous TF-target pairs compared to the best pair is less dramatic than in the mammalian species. Thus, the relative simplicity of the fly developmental system allows for the discovery of general trends, which are obscured, and thus more difficult to detect, in more complex (mammalian) systems.

Finally, further analysis at single cell level will contribute to understanding the temporal and spatial transcriptome determinants of cellular differentiation.

## Conclusions

In summary, we investigated the regulatory transcriptional programs underlying cell linages differentiation. Our results show that transcriptional changes occurring during cellular differentiation are likely to predominantly reflect the progressive loss of pluripotency and the gain of a mature metabolic state, as cells transition from undifferentiated to terminally differentiated states. These changes are common to most cell types, and dominate those underlying lineage specification and tissue specificity, which affect a relatively small number of genes. A network of TFs regulates, mainly, the cell differentiation transcriptional program and this GRN is conserved across metazoans during development. This novel gene regulatory program will help to better understanding of many differentiation events, both, *in vivo* and *in* vitro, and could contribute to the improvement of differentiation *in vitro* processes.

## Methods

### *Drosophila melanogaster* strains

Fly strains used for this study were: *nubbinGAL4*;UAS-GFP (selection of wing primordial cells), *GMRGAL4*;UAS-GFP (selection of eye cells) and *p{GAW}rnGAL4-5*;UAS-GFP (selection of leg cells and antenna cells), *enGAL4;UAS-GFP* (selection of anterior and posterior compartments in wing), *apGAL4;UAS-GFP* (selection of ventral and dorsal compartments in wing), *ubiRFP;p{GawB}C68a;UAS-GFP* (male and female genitalia discs). Flies were grown in standard media at 25ºC.

### Cell sorting, RNA isolation, library preparation and sequencing

Imaginal tissues from third instar larvae (110-115h after egg laying), early-pupa (120-130h) and late pharate (225-235h) were dissected in PBS 1x and incubated for 1h in a 10x trypsin solution (Sigma T4174) at room temperature in a rotating wheel. Cells were vigorously pipetted and kept on ice in Schneider’s insect medium. To discard dead cells, DAPI was added to the sample at 1 μg/mL final concentration. Cells were sorted in a FACSAria (BD) with the 85 μm nozzle at the Flow Cytometry Unit of the University Pompeu Fabra and the Centre for Genomic Regulation (UPF-CRG, Barcelona, Spain). Cells of interest were collected for subsequent analyses (**Supplementary Fig. 1B**). RNA from sorted cells was extracted with the ZR-RNA MicroPrep Kit from Zymo Research following the manufacturer’s instructions. Sequencing libraries were prepared using TruSeq Stranded mRNA Library Preparation Kit from Illumina and following the manufacturer’s instructions. Sequencing was performed in a HiSeq sequencer from Illumina at the Ultrasequencing Unit of the CRG. A minimum of 50 million paired-end 75 bp-long reads were obtained per replicate and two replicates were performed per each tissue.

### RNA-seq experiments

Stranded paired-end RNA-seq data for 27 samples in two bio-replicates were generated. The raw data (FASTQ), mapped data (BAM) and lists of quantified elements are available https://rnamaps.crg.cat/.

### RNA-seq data processing and analysis

We processed the data using the in-house pipeline grape-nf (available at https://github.com/guigolab/grape-nf). RNA-seq reads were aligned to the fly genome (dm6) using STAR 2.4.0j(58) allowing up to 4 mismatches per paired alignment. We used the FlyBase genome annotation r6.05(59,60). Only alignments for reads mapping to ten or fewer loci were considered. On average 92% of reads were mapped and 85% of the initial number of reads were uniquely mapped to the fly dm6 genome. Of these, 92% mapped to exonic regions.

Gene and transcripts were quantified in Transcripts Per Kilobase Million (TPMs) using RSEM (61). TPM values were recomputed including only protein coding and long non coding genes (13,920 and 2,470 genes, respectively). Only genes expressed at least 5 TPMs in two samples were considered for subsequent analyses (9334 genes). TF annotation was obtained from FlyFactorSurvey (http://mccb.umassmed.edu/ffs). Plots were made using d3js (available at https://d3js.org/) and ggplot2(62) and R scripts (some available at https://github.com/abreschi/Rscripts).

### Gene expression

#### Variance Decomposition

For each gene, the total variance in expression across samples (total sum of squares, TSSg) can be decomposed into three variances : variance across developmental stages (SSSg), variance across tissues (SSTg), and the residual variance (SSRg) as in the ANOVA type of analysis: TSSg=SSSg+SSTg+SSSg:SSTg+SSRg(19) The relative contribution of each factor to the total variance in gene expression can then be computed as the relative proportion of each variance with respect to the total. We used a linear model, implemented using the function lm() from R using the in-house wrapper available at https://github.com/abreschi/Rscripts/blob/master/anova.R (19). The TPM matrix with both replicates per sample was used for the analysis.

### Profiling the gene expression and DDGs definition

To identify genes whose expression changes across fly differentiation we used different methods to profile gene expression. First, we performed differential gene expression analysis using EdgeR v3.22.5 (63) with stages and tissues as factors. The counts matrix with both replicates per sample was used for the analysis. We required log2 fold change > 1 (at least two-fold change) and FDR < 0.01. Contrasts of every tissue and stage were used to define tissue and stage specific genes (18 contrast in total, herein called EdgeR tissue-stage). Second, to identify tissue-specific genes, EdgeR was used with tissue as factor (tissue gene profiles) and time-specific genes were classified for each tissue independently based on their trajectories (stage gene profiles: up-regulation, down-regulation, peaking or bending). Briefly, we focused on profiles with at least two-fold change and identified monotonic up-regulations and down-regulations; peaking profiles were defined as monotonic increases followed by monotonic decreases, bending profiles as the opposite (script: classification.log2.pl) (64). Third, we used the percentage of contribution from variance decomposition to identify genes for which the sum of tissue and stage contribution explains at least 70% of variation of expression.

Finally, genes classified as differentially expressed in at least two of the three methods used (edgeR tissue-stage, gene profiles, either across tissues or stages, and variance decomposition) were considered Developmental Dynamic Genes (DDGs).

In detail, EdgeR tissue-stage results were used to classify all gene sets: stage genes (differentially expressed in all the tissues in a particular stage/s), tissue genes (differentially expressed in all stages in a particular tissue/s and tissue-stage (differentially expressed in particular tissue and stage). Gene profiles through developmental stages were used to define stage genes and tissue-stage genes, while gene profiles across tissues were used to define tissue genes. Following variance decomposition classification: stage genes have stage-explained variation at least two fold greater than tissue-explained variation, tissue genes have tissue-explained variation at least two fold greater than stage-explained variation and tissue-stage were the rest of genes above the 70% cut-off. Groups with less than 5 genes were discarded. **Supplementary Table 6** summarizes the number of genes obtained from each analysis.

### Restricted vs widespread gene classification

Restricted genes show expression levels equal or higher than 5 TPMs only in the precise tissue/developmental stage where they are considered differentially expressed. The remaining differentially expressed genes are classified as widespread.

### GO term enrichment analysis

The GO term enrichment analysis for biological processes hierarchy was performed separately for each set of genes, with respect to all DDGs used as background. The enrichment is tested with the hypergeometric test implemented in the R package GOstats v2.44.0 (65). FlyBase gene IDs are converted to entrez gene IDs via the R package org.Dm.eg.db v3.4.1 (Bioconductor-org.Dm.eg.db), and mapped to gene ontology through the R package GO.db v3.4.1 (Bioconductor-GO.db).

### Epigenetic regulation

#### FAIRE-Seq data processing and classification

For each gene, we define the promoter as the sequence within a window of 250 bp upstream and downstream from the transcription start site (TSS). All TSSes annotated for the genes were considered for classification, but only the genes with all TSSes equally classified were considered for later analyses. FAIRE-Seq data was obtained from NCBI GEO database GSE38727 (21), replicates with higher signal-to-noise ratio were selected for the analysis. Data was processed using the in-house chip-nf pipeline (https://github.com/guigolab/chip-nf). Reads were continuously mapped to the fly genome (dm6) with up to two mismatches using the GEM mapper (66). Only alignments for reads mapping to 10 or fewer loci were reported. Duplicated reads were removed using Picard (http://broadinstitute.github.io/picard/). Peak calling was performed using MACS2 (67), only peaks with fdr < 0.1 were considered. Promoters were classified as specific when open chromatin peaks overlapping the promoter are present only in specific tissue and/or stage, as close when no open chromatin peaks overlap the promoter and as non specific when overlapping peaks are present in several tissues or stages.

#### H3K4me3 Chromatin Immunoprecipitation

Chromatin from antenna, eye, leg and wing at three stages of development (L3, EP and LP) was fixed with FA1% at RT for 10 min and sonicated with a Diagenode Bioruptor for 15 minutes at high intensity with ON/OFF alternate pulses of 30 second. Sheared chromatin was aliquoted and flash frozen in liquid nitrogen. Chromatin immunoprecipitation assays were performed following iChIP(68) protocol with some modifications. Abcam antibody Abcam_ab8895 was used to immunoprecipitate H3K4me3 attached chromatin. Data was processed using the in-house chip-nf pipeline (https://github.com/guigolab/chip-nf). Reads were continuously mapped to the fly genome (dm6) with up to two mismatches using the GEM mapper (66). Only alignments for reads mapping to 10 or fewer loci were reported. Duplicated reads were removed using Picard (http://broadinstitute.github.io/picard/). Peak calling was performed using MACS2 (67), only peaks with fdr < 0.1 were considered. The intersection of peaks called in both replicates were used for the analysis. The intersection of such peaks with accessible promoters, described in section “FAIRE-Seq data processing and classification” was used for the epigenetic analysis of promoter regions. The intersection of peaks was performed using BEDtools (69) intersectBed v2.17.0.

### Gene Regulatory network

#### Promoter open chromatin and motif search

As previous studies demonstrate that FAIRE-enriched regions(21,70,71) are bound by multiple regulatory factors, we used FAIRE-Seq data of fly from the same tissues and developmental stages we used for our analyses to identified putative regulators of DDGs. FAIRE-enriched regions overlapping DDGs promoters (window of 250 bp upstream and downstream from the transcription start site (TSS)) and open at least in the respective tissue and/or developmental stage where the gene is differentially expressed were selected for TF binding motif search. In the case of eye late pupa, for which data was not available, peak should be present in eye or CNS at L3 or in late pupa in any other tissue. The overlap between FAIRE-enriched regions and promoters was computed using BEDTools intersectBed v2.17.0 (69). We ran FIMO (72) using all available fly transcription factor (TF) motif matrices from MEME suite (73) against the FAIRE-enriched regions on DDGs promoters. We found conserved motifs for 238 fly TFs in the promoters of 1991 DDGs. Only TFs expressed at least 5 TPMs in two of the experiments were kept for building the network.

#### Sequence conservation and experimental data filtering

To predict binding sites in DDGs promoters every motif was inspected for conservation using the dm6 27-way multiple alignment (23 *Drosophila* species} sequences, house fly, *Anopheles* mosquito, honey bee and red flour beetle) and the phastCons measurement of evolutionary conservation from the UCSC Genome Browser (74,75). PhastCons scores in this window are averaged from the bigwig file with the bwtool software(76) and this average is taken as a measure of promoter sequence conservation. Alignment coverage should be at least 80% of initial fly input sequence in at least 10 species (at least one species further than *D.pseudoobscura*). Average phastCons over the motif region should be greater than 0.5.

#### Gene co-expression regulatory network

To build the fly gene co-expression regulatory network (GRN), we computed the correlation of expression between DDGs and potential regulatory TFs across all the samples produced here (including those from the wing compartments, the antenna and the genitalia). To generate GRN for DDGs, the R package WGCNA was used (27). We used expression values of DDGs and 775 fly TFs across the 27 samples generated in this study. Using default parameters of WGCNA package, that is: hierarchical clustering (hclust R function) and Dynamic tree cut R package (77) we identified fifteen clusters of expression (clusters with Pearson’s correlation coefficient higher than 0.85 were merged), and interactions were filtered first by coefficient of correlation (connection weight higher than 0.1) and then, by presence of a TF conserved motif in the accessible TSS promoter of the DDG. Software Cytoscape 3.8.0 was used for network visualization (78). Nodes were displayed according to Edge-weighted Spring-Embedded Layout analysis of TF-target correlation of expression (Pearson’s correlation coefficient between TF and target expression, averaged between replicates, calculated across the 27 samples) or the odes Betweenness centrality (measure of the amount of influence a node has over the flow of information in a graph calculated as the number of times a node acts as a bridge along the shortest path between two other nodes). Node size was adjusted depending on node closeness centrality (average shortest path length from the node to every other node in the network, it indicates how close a node is to all other nodes). Edges were colored according to TF-target correlation of expression (Pearson’s correlation coefficient mentioned above). Edge transparency was adjusted depending on TF-target weight (similarity measure considering levels of expression, averaged between replicates, across the 27 samples used for network generation).

#### Conservation of differentiation regulatory program in metazoans

Fly gene identifiers were mapped to mouse, human, and worm orthologs using Ensembl79 (http://mar2015.archive.ensembl.org) (79). Genes mapping to one or more orthologs in each species were analyzed in a fly-oriented manner.

Transcriptional profiling of mouse and human were obtained from ArrayExpress (E-MTAB-6798, and E-MTAB-6814) (24). Worm data was obtained from NCBI BioProject database (PRJNA477006) (32). The profile of gene expression of orthologs of *Drosophila* early and late genes along tissue development was analyzed using self-organizing maps (R package kohonen v3.0.8) (80,81). Orthologs that mapped into both early and late fly genes were excluded from all analyses. We compared the profiles of gene expression of fly early and late genes with the mouse orthologs based on the gene profile classification provided by the authors (24).

## Declarations

### Ethics approval and consent to participate

Not applicable

### Consent for publication

Not applicable

### Availability of data and materials

The datasets generated and analyzed during the current study from fly are available in the Array Express (https://www.ebi.ac.uk/arrayexpress/) repository, under accession number E-MTAB-10879 (for RNA-Seq samples) (http://www.ebi.ac.uk/arrayexpress/experiments/E-MTAB-10879); and under accession number E-MTAB-11307 (https://www.ebi.ac.uk/arrayexpress/experiments/E-MTAB-11307) (for ChIP-Seq data).

All the fly generated data is also available through the RNAmaps dashboard (https://rnamaps.crg.cat/).

The datasets analyzed during the current study from fly modENCODE are available in modENCODE portal repository (www.modencode.org) (28,30).

The datasets analyzed during the current study from fly midgut cell types is available in NCBI GEO repository under accession number GSE61361 (31).

The datasets analyzed during the current study from fly FAIRE-Seq data is available in NCBI GEO repository under accession number GSE38727 (21).

The datasets analyzed during the current study from mouse and human development are available in the from ArrayExpress (https://www.ebi.ac.uk/arrayexpress/) repository, under accession number E-MTAB-6798 and E-MTAB-6814 (24).

The dataset analyzed during the current study from worm development is available in the NCBI BioProject database repository under accession number PRJNA477006 (32).

### Author’s contributions

R.G. conceived the project. M.R-R. and R.G designed the study. C.C.K, M.R-R. and A.B. performed the computational analyses. S.P-L., M.R-R. and A.A. performed the RNA-Seq experiments. S.P-L. and M.R-R. performed the ChIP-Seq experiments. M.R-R., C.C.K and R.G. wrote the manuscript with the contribution of all authors. All authors read and approved the final manuscript.

## Acknowledgements

We thank Emilio Palumbo for his help with pipeline development and data processing, Bruna R. Correa, Ramil Nurtdinov and Beatrice Borsari for helpful discussions about the data and the manuscript and Romina Garrido for administrative assistance. We also thank the CRG Genomics Unit and the CRG/UPF Flow Cytometry Unit (Barcelona, Spain).

