## supplementary material for "A transcriptional program shared across lineages underlies cell differentiation during metazoan development"

**This PDF file includes:**

Figs. S1 to S11

Tables S1, S4 and S6

**Other Supplementary information for this manuscript include the following:**

Tables S2, S3 and S5

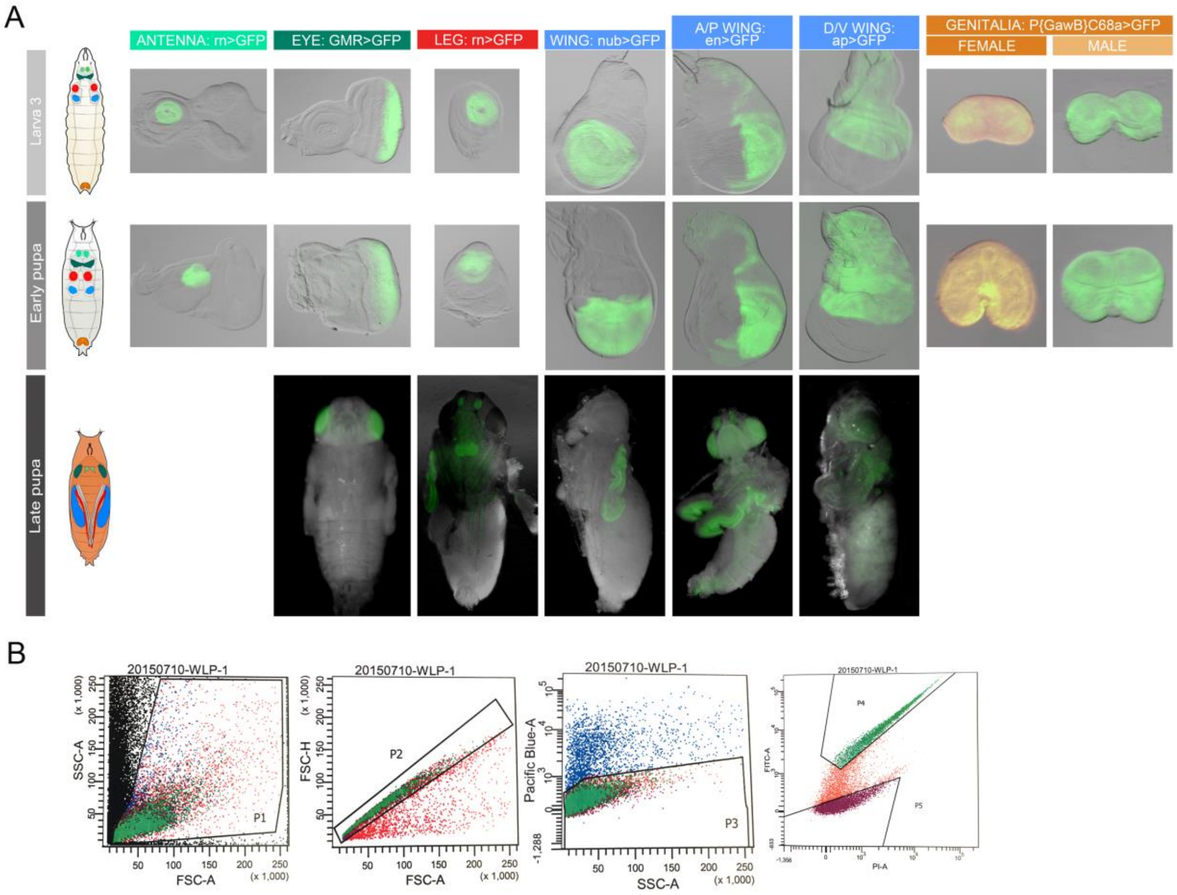


### Supplementary Fig. 1.

***Drosophila melanogaster*** **imaginal discs cell isolation and generation of the data**. **(A)** Images of antenna, eye, leg, wing and genitalia tissues at third instar larva, early pupa and late pupa. For each tissue we used a particular promoter driving GFP expression (green fluorescence signal in each picture) in the cells of interest. **(B)** FACS analysis in a wing late pupa (WLP) sample. SSC-A (side scatter area), FSC (forward scatter), SSC-H (side scatter height), Pacific Blue and FITC (correspond to dye emission signaling). P1: integer cells, P2: single cells, P3: alive cells, P4: GFP positive cells, corresponding to wing blade cells. P4 population (GFP positive) was collected for wing RNA-Seq.

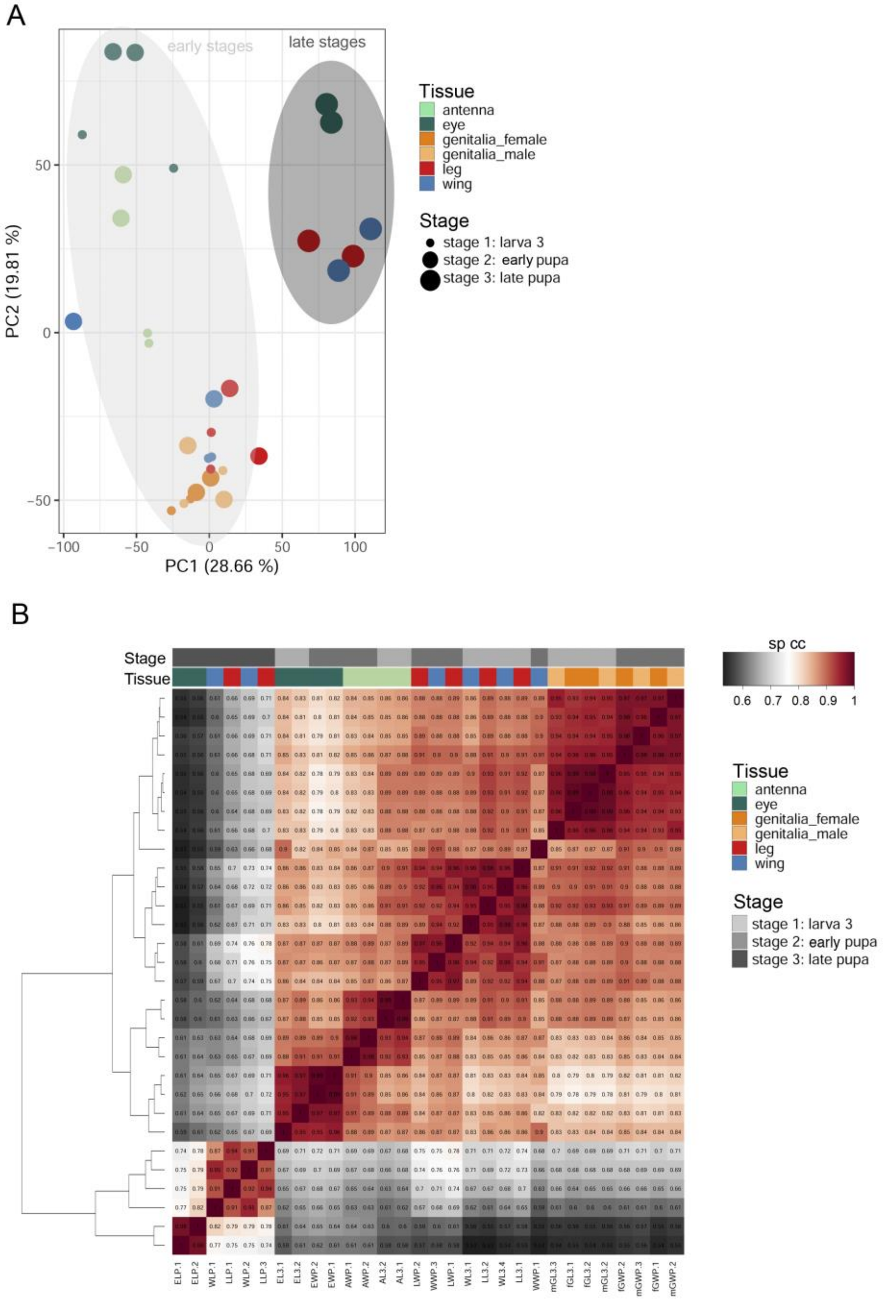


### Supplementary Fig. 2.

**Clustering of RNA-Seq samples**. **(A)** Principal component analysis (PCA) based on the expression of all expressed genes (with at least 5 TPMs in at least two samples).

PC1 separates early and late stages. PC2 separates neural tissues (antenna and eye) from the non-neural ones. **(B)** Clustering of RNA-Seq samples based on the Spearman's correlation coefficient (sp cc) of gene expression of all expressed genes (with at least 5 TPMs in at least two samples).

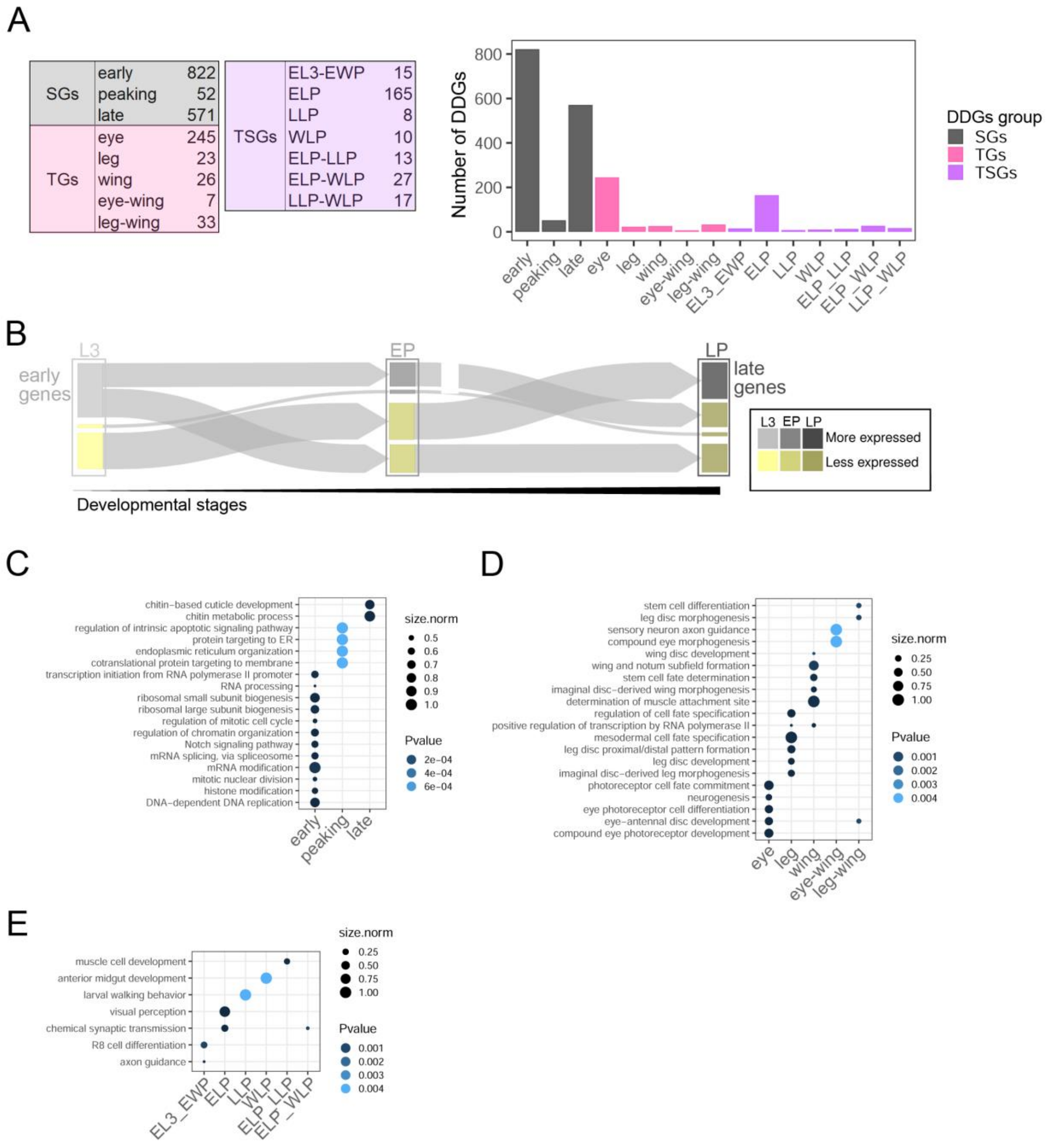


**Supplementary Fig. 3.**

**Classification, expression dynamics and functional characterization of DDGs. (A)** Number of DDGs in the different expression categories. **(B)** Dynamics of genes differentially expressed across developmental stages (SGs) represented as a Sankey diagram. We represent the genes that are more expressed (upper) and less expressed (lower) at each developmental stage. The arrows represent the number of genes that transition from one development stage to the next from differentially expressed to less expressed (or vice versa). About half of the early genes are specifically expressed at L3, while the other half are also expressed at EP. **(C)** Gene Ontology (GO) term enrichment analysis of SGs. The dot size represents the normalized number of genes in each category. **(D)** GO term enrichment analysis of TGs. The dot size represents the normalized number of genes in each category. **(E)** GO term enrichment analysis of TSGs. The dot size represents the normalized number of genes in each category.

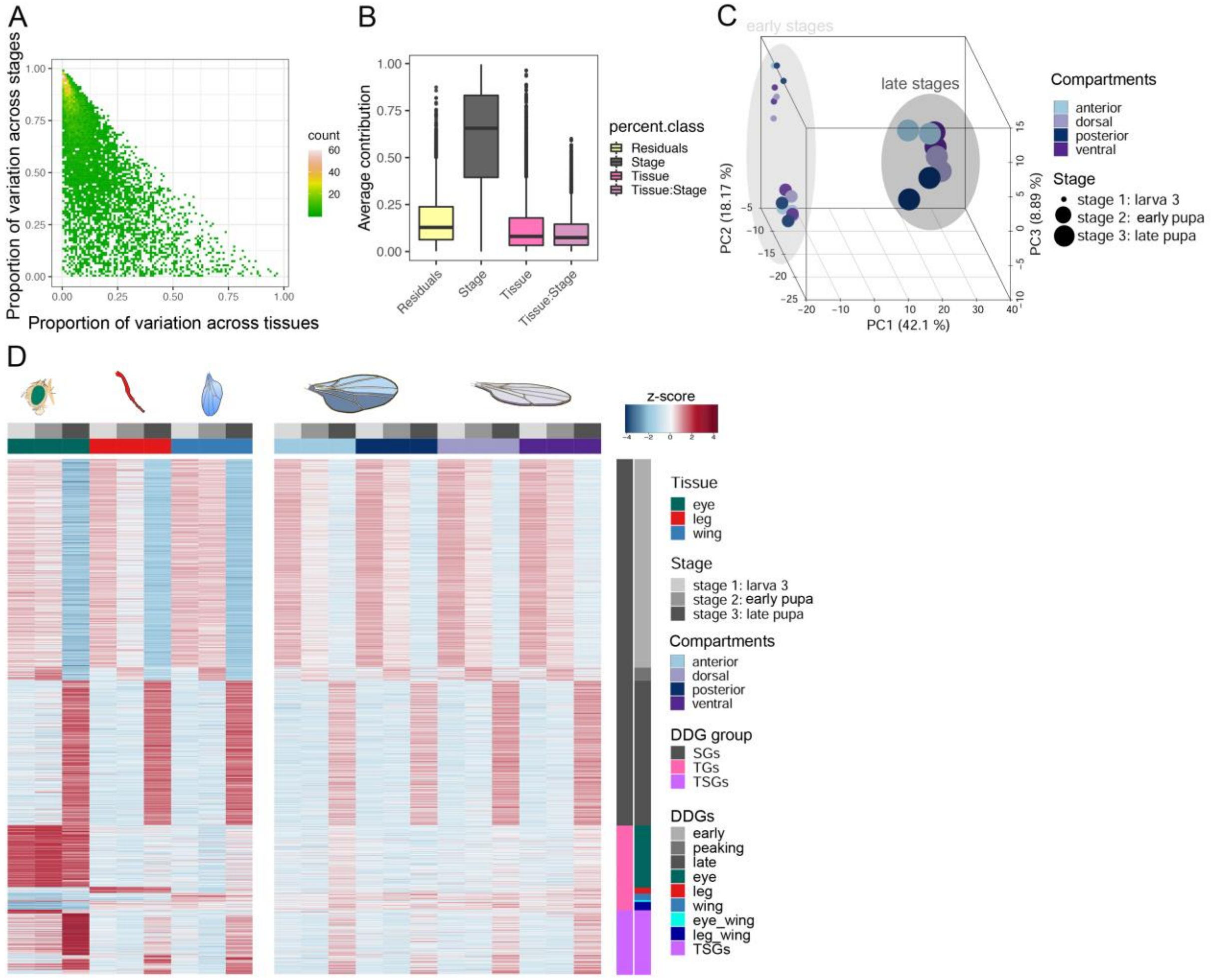


### Supplementary Fig. 4.

**Gene expression dynamics along fly development in wing compartments. (A)** Proportion of the variance in gene expression explained by wing compartments (x-axis) and by developmental stages (y-axis). Each dot corresponds to one of the 7,683 genes that are expressed at least 5 TPMs in at least two samples. **(B)** Proportion of gene expression variance explained by stage, tissue and the interaction between the two. **(C)** Principal component analysis (PCA) based on the expression of the 1,000 most variable genes across tissues, compartments and developmental stages (same set of genes as in Fig. 1C). PC1 separates the early and the late stages. PC2 separates compartments in the late stage. PC3 separates L3 and EP stages. **(D)** DDGs expression across tissues (left, as Fig. 2C) and compartments (right). Gene expression values are normalized to z-score values.

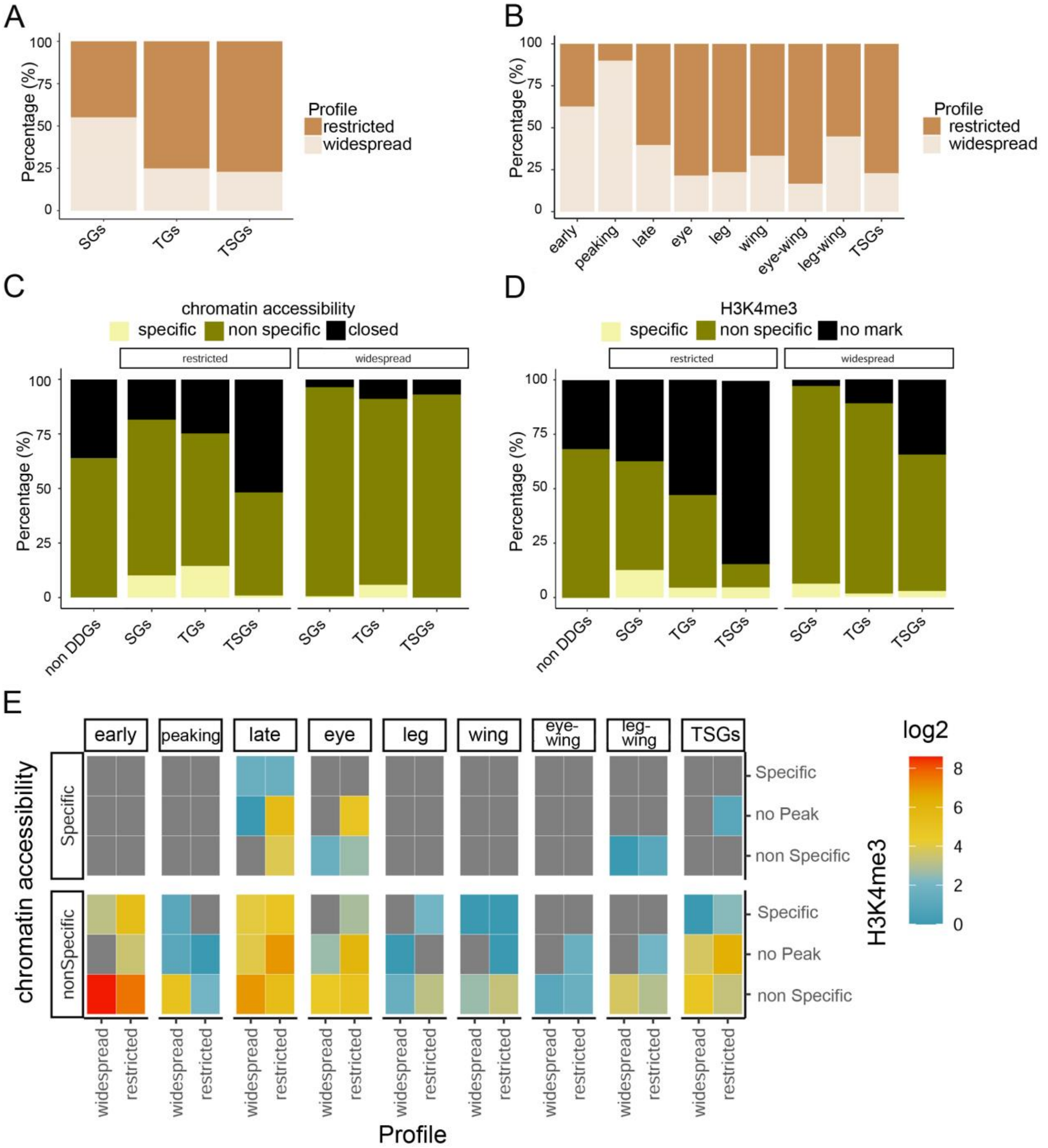


**Supplementary Fig. 5.**

**Epigenetic regulation of DDGs. (A** and **B)** Percentage of DDGs with restricted/widespread expression. **(C)** Chromatin accessibility based on FAIRE-Seq at the promoters of DDGs. Promoters are classified as closed (when there is no FAIRE-Seq peak in any time point and/or tissue), specific (when a peak is present exclusively in the time point and/or tissue where the gene is differentially expressed) and non specific (when peak is present in several time points and/or tissues). **(D)** H3K4me3 marking at the promoters of DDGs. Promoters are classified as no mark (when there is no H3K4me3 peak in any time point and/or tissue), specific (when a peak is present exclusively in the time point and/or tissue where the gene is differentially expressed) and non specific (when a peak is present in several time points and/or tissues). **(E)** Total number of genes in each DDG category depending on developmental expression pattern, specificity of chromatin accessibility and H3K4me3 and breadth of expression.

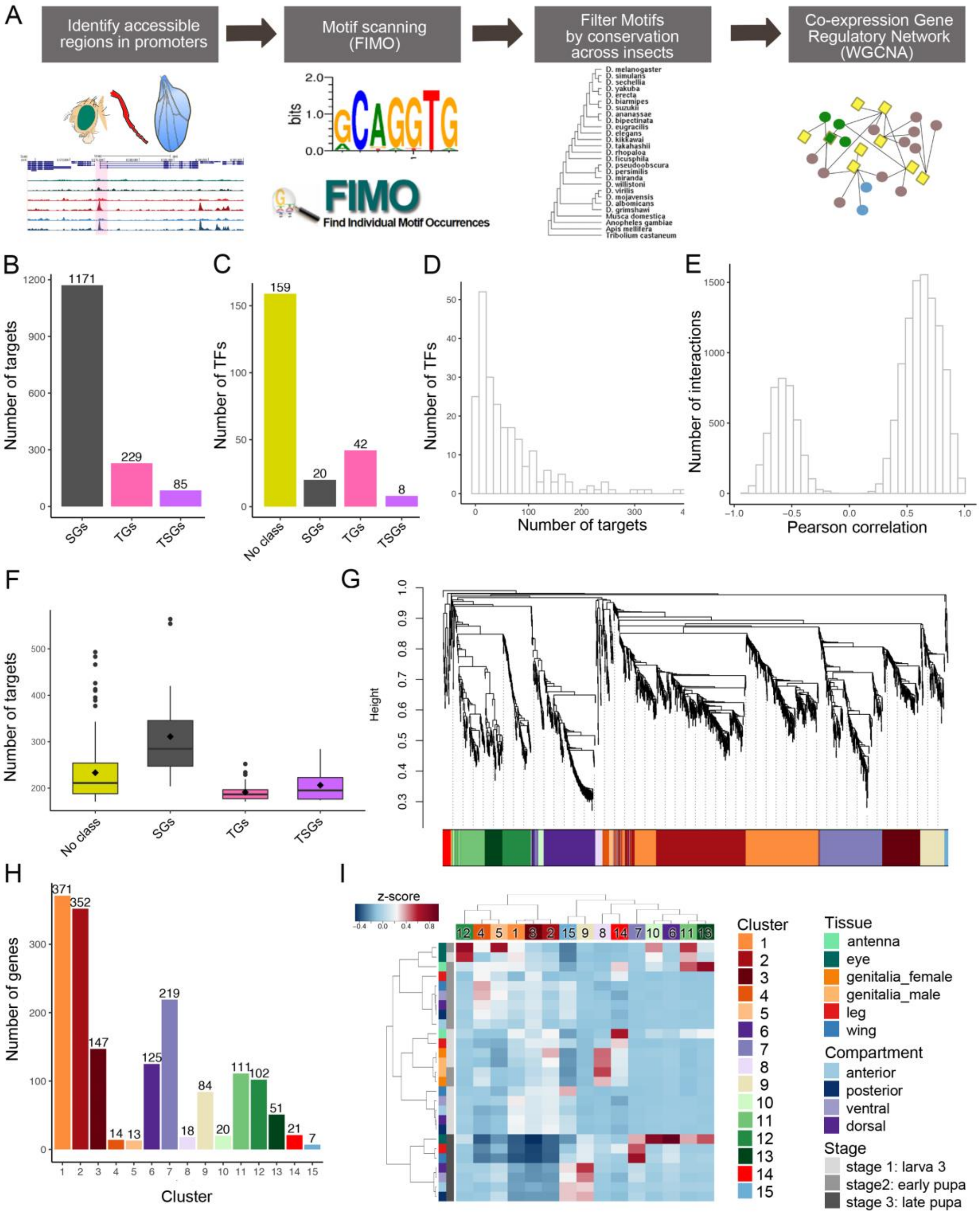


**Supplementary Fig. 6.**

**Construction and properties of the *Drosophila* gene regulatory network (GRN) (A)** Pipeline to generate the GRN (see Methods for details). **(B)** Number of target genes in the network belonging to the different DDG categories (total number, 1,485). **(C)** Number of TFs in the network in each DDG category (total number, 229). **(D)** Number of targets per TF. **(E)** Distribution of positive (Pearson’s correlation coefficient > 0.3) and negative (Pearson’s correlation coefficient < -0.3) interactions (14,039 in total). **(F)** Number of targets for TFs in each DDG category. **(G)** Hierarchical clustering based on expression of the genes in the GRN. The colored bar at the bottom indicates the network clusters identified by the WGCNA. Height corresponds to one minus Pearson’s correlation. **(H)** Number of genes in each cluster. **(I)** Cluster eigengene expression in tissues, compartments and developmental stages. Intuitively, the eigengene represents the archetypical gene within the cluster. For instance, the eigenexpression of cluster 9 (late genes) is indeed high in a number of late tissues. Gene expression values are normalized to z-score values.

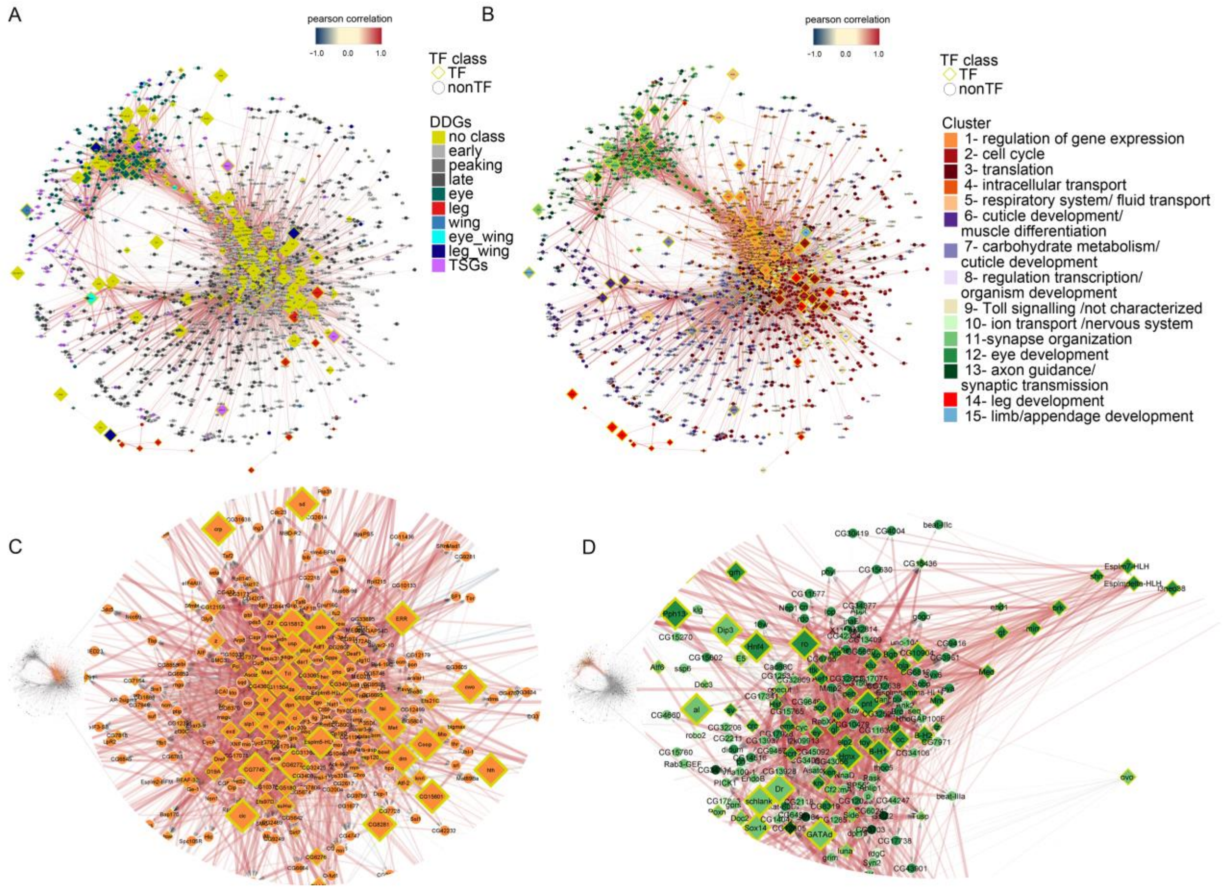


**Supplementary Fig. 7.**

***Drosophila* gene regulatory network (GRN)**. **(A)** GRN displayed according to node betweenness centrality (i.e. how central the node is in the graph, see Methods). Edges are colored according to TF-target Pearson's correlation coefficient (red > 0.3, blue < -0.3). Nodes are colored by DDG category. Node size reflects node closeness centrality. **(B)** GRN displayed according to node betweenness centrality. Nodes are colored by cluster. Note that Cluster 1 genes, that are associated with gene expression regulation, show higher degree of centrality compared to other genes. **(C)** Detailed view of most TFs from cluster 1. **(D)** Detailed view of TFs belonging to eye clusters 10, 11,12 and 13.

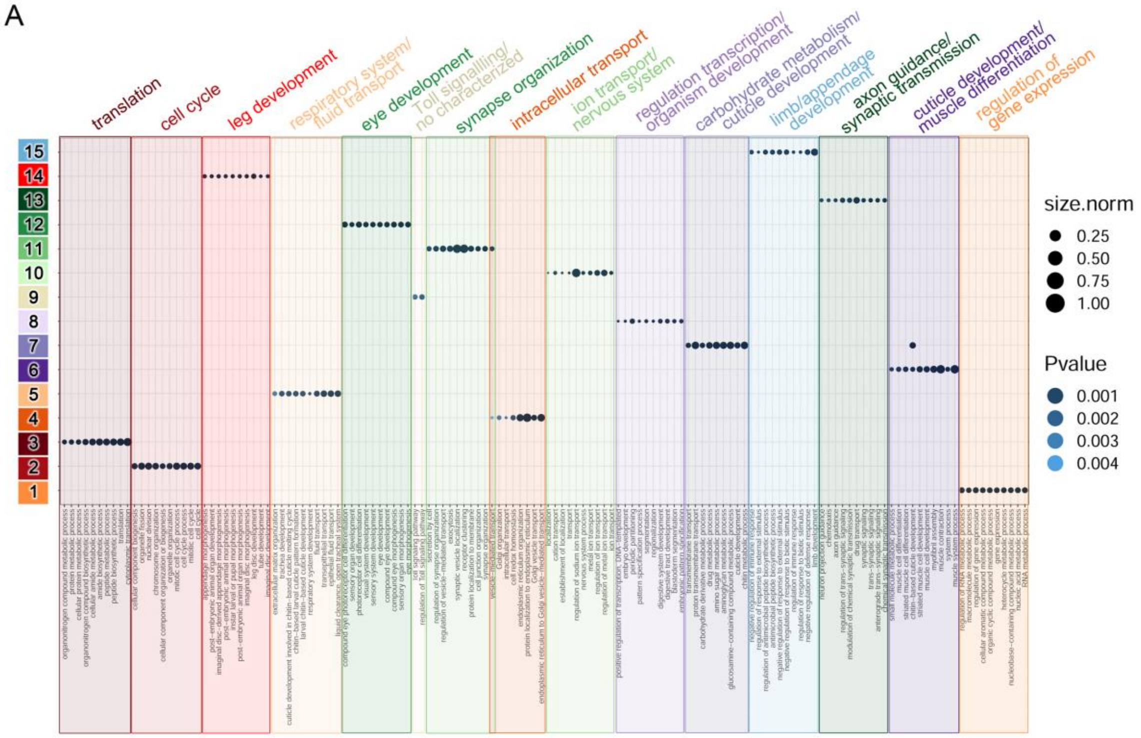


**Supplementary Fig. 8.**

**Functional characterization of the *Drosophila* gene regulatory network (GRN).** Gene Ontology enrichments in each cluster (the top 10 most significant functional categories are shown). Clusters tend to be associated with a particular functional category. The dot size represents the normalized number of genes in each category.

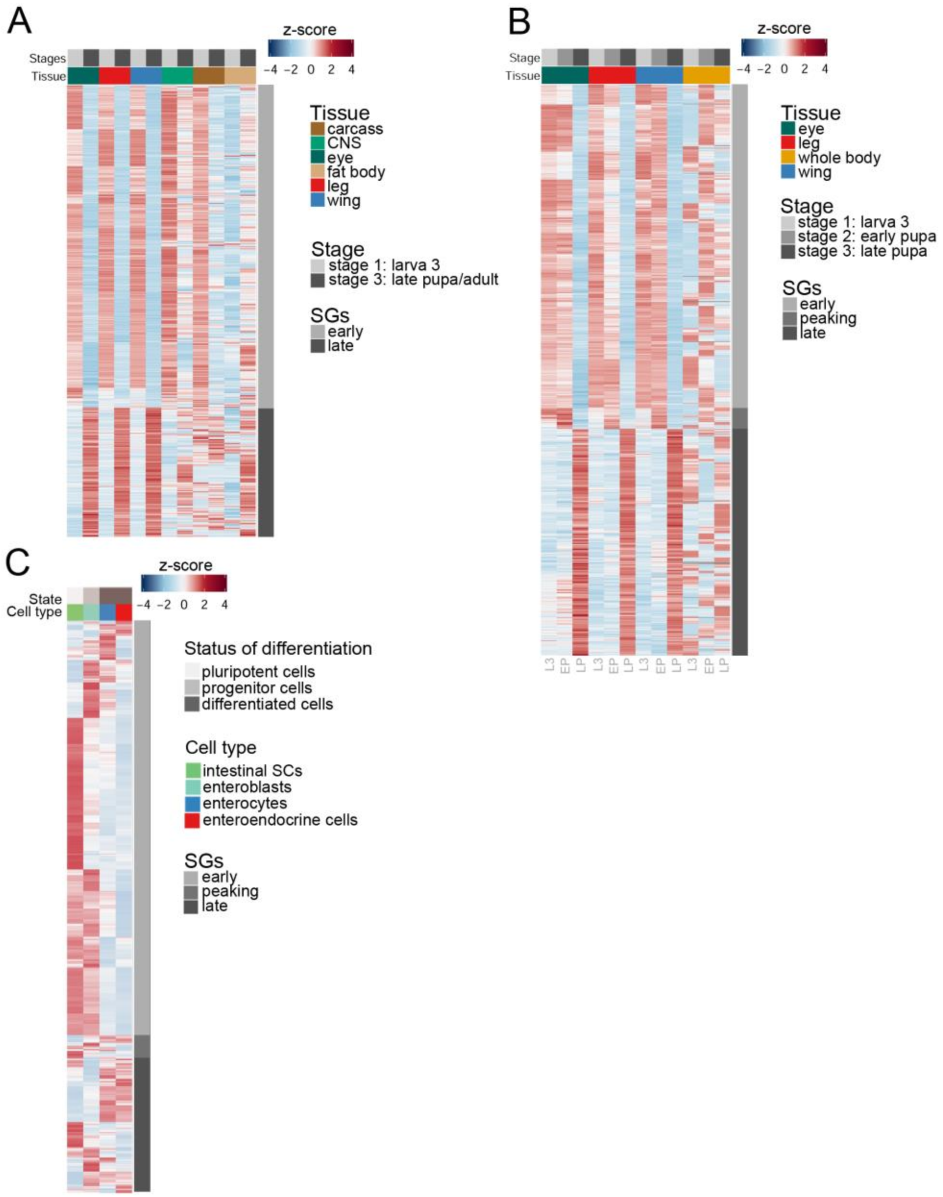


**Supplementary Fig. 9.**

**Gene Expression dynamics of SGs in tissues, whole animal and cell types during fly development and differentiation. (A)** Expression of SGs in modENCODE tissues and in the imaginal tissues monitored here at L3 and LP (or adult for carcass) stages. **(B)** Expression of SGs in modENCODE whole animals at L3, EP and LP stages (same data as in Fig 5b, but with the z-score computed considering only samples from these three time points. **(C)** Expression of SGs in midgut cell types: intestinal stem cells (light green), enteroblasts (progenitor cells, cian), enterocytes (blue) and enteroendocrine cells (red).

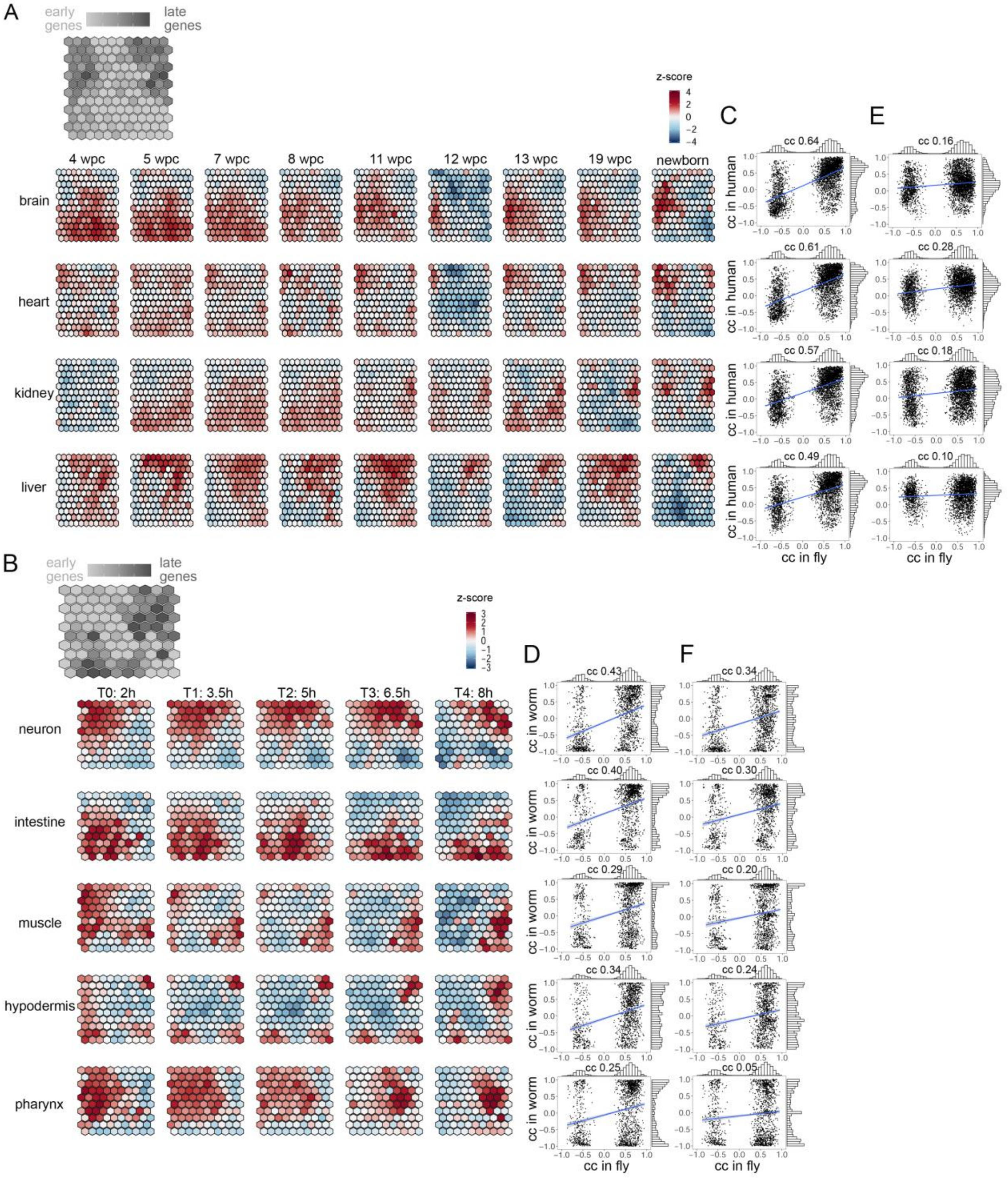


**Supplementary Fig. 10.**

**Conservation of the *Drosophila* GRN in human and worm**. **(A)** SOM of human orthologs of fly early and late genes based on their expression in a number of tissues during development (brain, heart, kidney and liver) at different weeks post conception (wpc) and newborn individuals. Early (light grey) and late (dark grey) orthologs cluster separately. **(B)** SOM of worm orthologs of fly early and late genes based on their expression in a number of tissues during development (neuron, intestine, muscle, hypodermis and pharynx) from 2h to 8h after embryo isolation. Early (light grey) and late (dark grey) orthologs cluster separately. **(C)** Scatter plots of the correlation in fly and in human of TF-target pair. The correlations have been computed independently in each fly-human ortholog tissue. When multiple human orthologous targets are found for the same orthologous TF, the TF-target pair with the closest correlation is employed. Pearson's correlation coefficient (cc) between fly and human correlations is shown on the top of the plots, p-values of the correlations are 0 for brain, heart and kidney and $1.09^{e-317}$ for liver. **(D)** Scatter plots of the correlation in fly and in worm of TF-target pair. The correlations have been computed independently in each fly-human ortholog tissue. When multiple worm orthologous targets are found for the same orthologous TF, the TF-target pair with the closest correlation is employed. Pearson's correlation coefficient (cc) between fly and worm correlations is shown on the top of the plots, p-values of the correlations are $1.45^{e-75}$, $7.50^{e-58}$, $4.02^{e-32}$, $8.13^{e-41}$ and $2.43^{e-25}$ for neuron, intestine, muscle, hypodermis and pharynx, respectively. **(E)** Scatter plots showing the same correlation as in (C) panel but considering all possible orthologous pairs, when multiple human orthologous targets are found for the same orthologous TF, the mean of the correlations is employed. Pearson's correlation coefficient (cc) between fly and mouse correlations is shown on the top of the plots, p-values of the correlations are $7.71^{e-23}$, $1.05^{e-70}$, $9.73^{e-32}$ and $1.30^{e-08}$ for brain, heart, kidney and liver, respectively. **(F)** Scatter plots showing the same correlation as (D) panel but considering all possible orthologous pairs. When multiple human orthologous targets are found for the same orthologous TF, the mean of the correlations is employed. Pearson's correlation coefficient (cc) between fly and worm correlations is shown on the top of the plots, p-values of the correlations are $6.78^{e-47}$, $4.21^{e-31}$, $4.10^{e-16}$, $5.79^{e-20}$ and $1.08^{e-05}$ for neuron, intestine, muscle, hypodermis and pharynx, respectively.

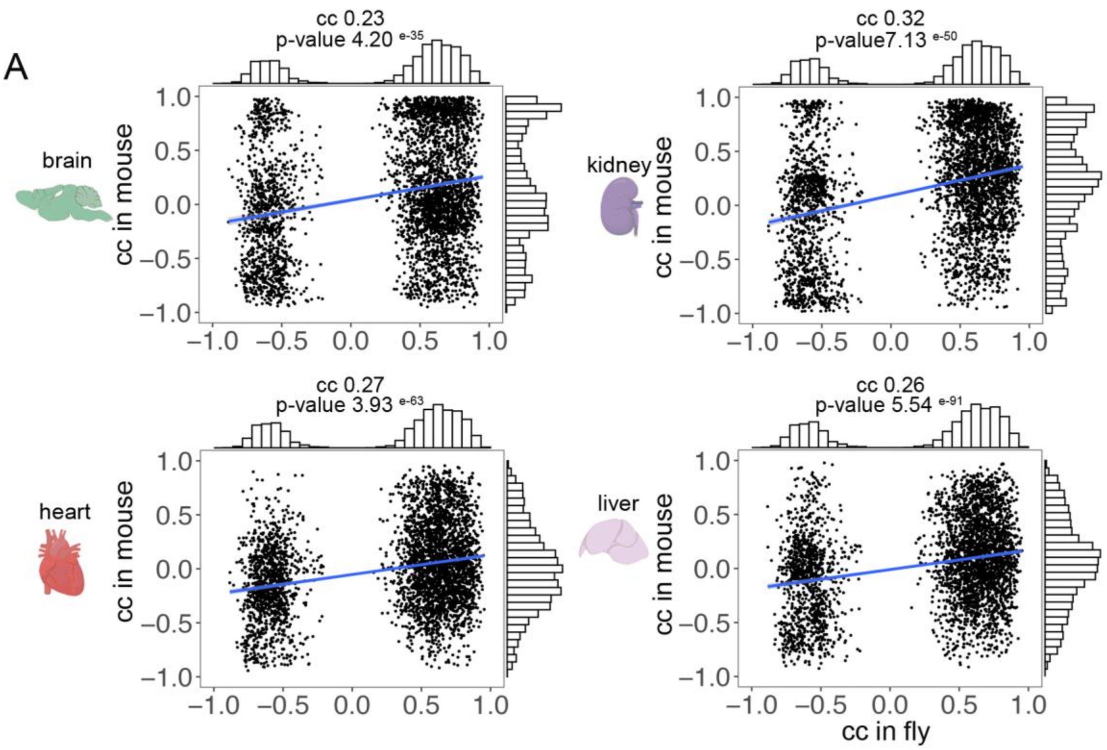


**Supplementary Fig. 11.**

**Conservation in mouse of the fly GRN TF-target associations**. Pearson's correlation between the expression of the mouse orthologs for the fly GRN TF-target pairs. In contrast to Fig. 6F, here, correlations are computed across all TF-target mouse orthologous pairs. When multiple mouse orthologous targets are found for the same orthologous TF, the mean of the correlations is employed. Pearson's correlation coefficient (cc) between fly and mouse correlations and p-values of the correlations are shown on the top of the plots. (Figure partially created with BioRender.com).

**Supplementary Table 1.**

**Control genes.** List of genes known to be differentially expressed between eye leg and wing or L3, EP and LP.

| Flybase_id | gene_name | control for |
|---|---|---|
| FBgn0260400 | *elav* | eye |
| FBgn0004595 | *pro* | eye |
| FBgn0003460 | *so* | eye |
| FBgn0002940 | *ninaE* | eye |
| FBgn0003366 | *sev* | eye |
| FBgn0000206 | *boss* | eye |
| FBgn0004618 | *gl* | eye |
| FBgn0003285 | *rst* | eye |
| FBgn0005558 | *ey* | eye |
| FBgn0000108 | *Appl* | eye |
| FBgn0000320 | *eya* | eye |
| FBgn0019650 | *toy* | eye |
| FBgn0028991 | *seq* | eye |
| FBgn0000157 | *dll* | leg |
| FBgn0003944 | *ubx* | leg |
| FBgn0025525 | *bab2* | leg |
| FBgn0026411 | *lim1* | leg |
| FBgn0001981 | *esg* | leg |
| FBgn0004870 | *bab1* | leg |
| FBgn0000061 | *al* | leg |
| FBgn0261648 | *salm* | wing |
| FBgn0001319 | *kn* | wing |
| FBgn0267337 | *rn* | wing |
| FBgn0004101 | *bs* | wing |
| FBgn0003975 | *vg* | wing |
| FBgn0004009 | *wg* | wing |
| FBgn0000492 | *Dr* | wing |
| FBgn0085424 | *nub* | wing |

| | | |
|---|---|---|
| FBgn0003984 | *vn* | wing |
| FBgn0086680 | *vvl* | wing |
| FBgn0002563 | *Lsp1beta* | L3 |
| FBgn0040718 | *CG15353* | L3 |
| FBgn0000055 | *Adh* | L3 |
| FBgn0002531 | *Lcp1* | L3 |
| FBgn0002533 | *Lcp2* | L3 |
| FBgn0002535 | *Lcp4* | L3 |
| FBgn0002789 | *Mp20* | L3 |
| FBgn0002564 | *Lsp1gamma* | L3 |
| FBgn0003721 | *Tm1* | L3 |
| FBgn0002773 | *Mlc2* | L3 |
| FBgn0005666 | *bt* | L3 |
| FBgn0002562 | *Lsp1alpha* | L3 |
| FBgn0004028 | *wupA* | L3 |
| FBgn0015664 | *Dref* | L3 |
| FBgn0011703 | *RnrL* | L3 |
| FBgn0024227 | *aurB* | L3 |
| FBgn0015271 | *Orc5* | L3 |
| FBgn0001086 | *fzy* | L3 |
| FBgn0263855 | *BubR1* | L3 |
| FBgn0050011 | *gem* | L3 |
| FBgn0000996 | *dup* | L3 |
| FBgn0000405 | *CycB* | L3 |
| FBgn0015278 | *Pi3K68D* | L3 |
| FBgn0037249 | *eIF3-S10* | L3 |
| FBgn0017577 | *Mcm5* | L3 |
| FBgn0015270 | *Orc2* | L3 |
| FBgn0038499 | *Brf* | L3 |
| FBgn0029840 | *raptor* | L3 |
| FBgn0250906 | *Pgk* | EP |
| FBgn0012036 | *Aldh* | EP |
| FBgn0259176 | *bun* | EP |
| FBgn0010548 | *Aldh-III* | EP |

| | | |
|---|---|---|
| FBgn0001091 | *Gapdh1* | EP |
| FBgn0003074 | *Pgi* | EP |
| FBgn0005619 | *Hdc* | EP |
| FBgn0001257 | *ImpL2* | EP |
| FBgn0260635 | *Diap1* | EP |
| FBgn0004108 | *Nrt* | EP |
| FBgn0011706 | *rpr* | EP |
| FBgn0267385 | *PyK* | EP |
| FBgn0004885 | *tok* | EP |
| FBgn0000064 | *Ald* | EP |
| FBgn0086355 | *Tpi* | EP |
| FBgn0010113 | *hdc* | EP |
| FBgn0025741 | *PlexA* | EP |
| FBgn0000382 | *csw* | EP |
| FBgn0010052 | *Jhe* | EP |
| FBgn0000551 | *Edg78E* | EP |
| FBgn0000552 | *Edg84A* | EP |
| FBgn0263782 | *Hmgcr* | EP |
| FBgn0039481 | *Cpr97Eb* | EP |
| FBgn0033359 | *CG8213* | LP |
| FBgn0003382 | *sha* | LP |
| FBgn0010611 | *Hmgs* | LP |
| FBgn0010470 | *Fkbp14* | LP |
| FBgn0085234 | *CG34205* | LP |
| FBgn0035398 | *Cht7* | LP |
| FBgn0004167 | *kst* | LP |
| FBgn0035512 | *Cpr64Ac* | LP |
| FBgn0066365 | *dyl* | LP |
| FBgn0005626 | *ple* | LP |
| FBgn0250833 | *CG34461* | LP |
| FBgn0013717 | *not* | LP |
| FBgn0001311 | *kkv* | LP |
| FBgn0001321 | *knk* | LP |
| FBgn0003525 | *stg* | LP |

| | | |
|---|---|---|
| FBgn0002577 | *m* | LP |
| FBgn0004511 | *dy* | LP |

**Supplementary Table 4.**

**Number of 1-to-many orthologs for early and late genes.** Number of early and late genes orthologs considered in the comparison between fly, mouse, human and worm.

| | mouse | human | worm |
|---|---|---|---|
| early | 1022 | 979 | 590 |
| late | 466 | 453 | 252 |

**Supplementary Table 6.**

**Total number of DDGs and number of genes classified using each method of gene profiling.** Classification of genes according to each method. Genes classified equally with at least two methods were considered DDGs.

| | DDGs classification | Variance Decomposition | tissue profile | time profile | edgeR |
|---|---|---|---|---|---|
| stage | 1445 | 2784 | | 1162 | 957 |
| tissue | 334 | 741 | 401 | | 210 |
| tissue-stage | 225 | 674 | | | 1091 |
| all | 2004 | 4199 | 401 | 1162 | 2258 |
